# The intra-genus and inter-species quorum-sensing autoinducers exert distinct control over *Vibrio cholerae* biofilm formation and dispersal

**DOI:** 10.1101/713685

**Authors:** Andrew A. Bridges, Bonnie L. Bassler

**Affiliations:** Department of Molecular Biology, Princeton University, Princeton, NJ 08544, USA; Howard Hughes Medical Institute, Chevy Chase, MD 20815, USA

## Abstract

*Vibrio cholerae* possesses multiple quorum-sensing systems that control virulence and biofilm formation among other traits. At low cell densities, when quorum-sensing autoinducers are absent, *V. cholerae* forms biofilms. At high cell densities, when autoinducers have accumulated, biofilm formation is repressed and dispersal occurs. Here, we focus on the roles of two well-characterized quorum-sensing autoinducers that function in parallel. One autoinducer, called CAI-1, is used to measure *vibrio* abundance, and the other autoinducer, called AI-2, is a broadly-made universal autoinducer that is presumed to enable *V. cholerae* to assess the total bacterial cell density of the vicinal community. The two *V. cholerae* autoinducers funnel information into a shared signal relay pathway. This feature of the quorum-sensing system architecture has made it difficult to understand how specific information can be extracted from each autoinducer, how the autoinducers might drive distinct output behaviors, and in turn, how the bacteria use quorum sensing to distinguish self from other in bacterial communities. We develop a live-cell biofilm formation and dispersal assay that allows examination of the individual and combined roles of the two autoinducers in controlling *V. cholerae* behavior. We show that the quorum-sensing system works as a coincidence detector in which both autoinducers must be present simultaneously for repression of biofilm formation to occur. Within that context, the CAI-1 quorum-sensing pathway is activated when only a few *V. cholerae* cells are present, whereas the AI-2 pathway is activated only at much higher cell density. The consequence of this asymmetry is that exogenous sources of AI-2, but not CAI-1, contribute to satisfying the coincidence detector to repress biofilm formation and promote dispersal. We propose that *V. cholerae* uses CAI-1 to verify that some of its kin are present before committing to the high-cell-density quorum-sensing mode, but it is, in fact, the universal autoinducer AI-2, that sets the pace of the *V. cholerae* quorum-sensing program. This first report of unique roles for the different *V. cholerae* autoinducers suggests that detection of self fosters a distinct outcome from detection of other.

## Introduction

Bacteria communicate and orchestrate collective behaviors using a process called quorum sensing (QS). QS relies on the production, release, and group-wide detection of extracellular signaling molecules called autoinducers. QS allows bacteria to assess the cell density and the species composition in the local environment and change their behavior accordingly [1,2]. Frequently, QS controls the development of biofilms, which are surface-associated communities of bacteria that secrete an adhesive extracellular matrix [3,4]. Biofilms are beneficial in many contexts, for example, the microbiota of the digestive tract exist in biofilms, but biofilms can also be harmful, for example, in infections. Cells in biofilms display striking differences from their planktonic counterparts, including extracellular matrix production and a dramatic tolerance to environmental perturbations, including antibiotic treatment [4,5]. Despite the extraordinary importance of bacterial biofilms, we know only a few key facts about their development: matrix production is required, and QS-mediated communication is involved in regulating biofilm formation and dispersal [4,6,7].

The pathogen and model QS bacterium *Vibrio cholerae* forms biofilms in all of its niches [4,8]. *V. cholerae* strains locked in the low-cell-density (LCD) QS mode avidly form biofilms, while strains locked in the high-cell-density (HCD) QS mode are incapable of forming biofilms [3]. While these findings show an overarching role for QS in repressing biofilm formation at HCD, they are incomplete because they were obtained from *V. cholerae* mutants locked in the LCD or HCD QS mode that are thus unable to progress through the normal QS program. Furthermore, how *V. cholerae* cells disperse from biofilms and the role played by QS in dispersal have only recently begun to be addressed [9]. Here, we establish a simple microscopy-based assay with wildtype (WT) *V. cholerae* that allows us to examine the full biofilm lifecycle and assess the role of QS in both biofilm formation and biofilm dispersal.

The canonical *V. cholerae* QS pathway is composed of two well-characterized autoinducer-receptor pairs that function in parallel to funnel cell-density information internally to control gene expression (Fig 1A) [10]. One autoinducer-receptor pair consists of *cholerae* autoinducer-1 (CAI-1; ((S)-3-hydroxytridecan-4-one)), produced by CqsA and detected by the two-component sensor-histidine kinase, CqsS [11,12]. CAI-1 is an intra-genus signal for *vibrios*. The second autoinducer-receptor pair is comprised of autoinducer-2 (AI-2; (2S,4S)-2-methyl-2,3,3,4-tetrahydroxytetrahydrofuran borate), produced by the broadly-conserved synthase, LuxS, and detected by LuxPQ [10,13,14]. LuxP is a periplasmic binding protein that interacts with AI-2. LuxP ligand occupancy is monitored by LuxQ, a transmembrane two-component sensor-histidine kinase [15,16]. AI-2 is produced by diverse bacterial species and is considered a universal QS autoinducer that conveys inter-species information [14]. Two other receptors, CqsR and VpsS, have recently been shown to feed information into this network, however the identities of their cognate autoinducers are not known (Fig 1B) [17,18]. Ethanolamine functions as a surrogate agonist for CqsR [19]. All four receptors act as kinases at LCD in their un-liganded states [20–22]. They funnel phosphate through the phospho-transfer protein LuxU to the response regulator LuxO, which, via a set of small regulatory RNAs (sRNAs) called the Qrr sRNAs, drives the production of the LCD master regulator AphA and represses production of the HCD master regulator HapR [23–26]. Under these conditions, behaviors including biofilm formation and virulence factor production are undertaken (Fig 1A, left) [3,10]. When bound to their cognate autoinducers, the receptors act as phosphatases [21]. LuxO is dephosphorylated, AphA production is terminated, and HapR production is activated [26]. In this situation, HCD behaviors are enacted, and, germane to this work, virulence and biofilm formation are repressed, and *V. cholerae* disperses from existing biofilms (Fig 1A, right) [9,27]. It has long been puzzling why the autoinducer signals are processed via the identical, shared pathway in *V. cholerae* as this system architecture is not obviously conducive to gleaning specific information from each autoinducer.

**Fig 1.**
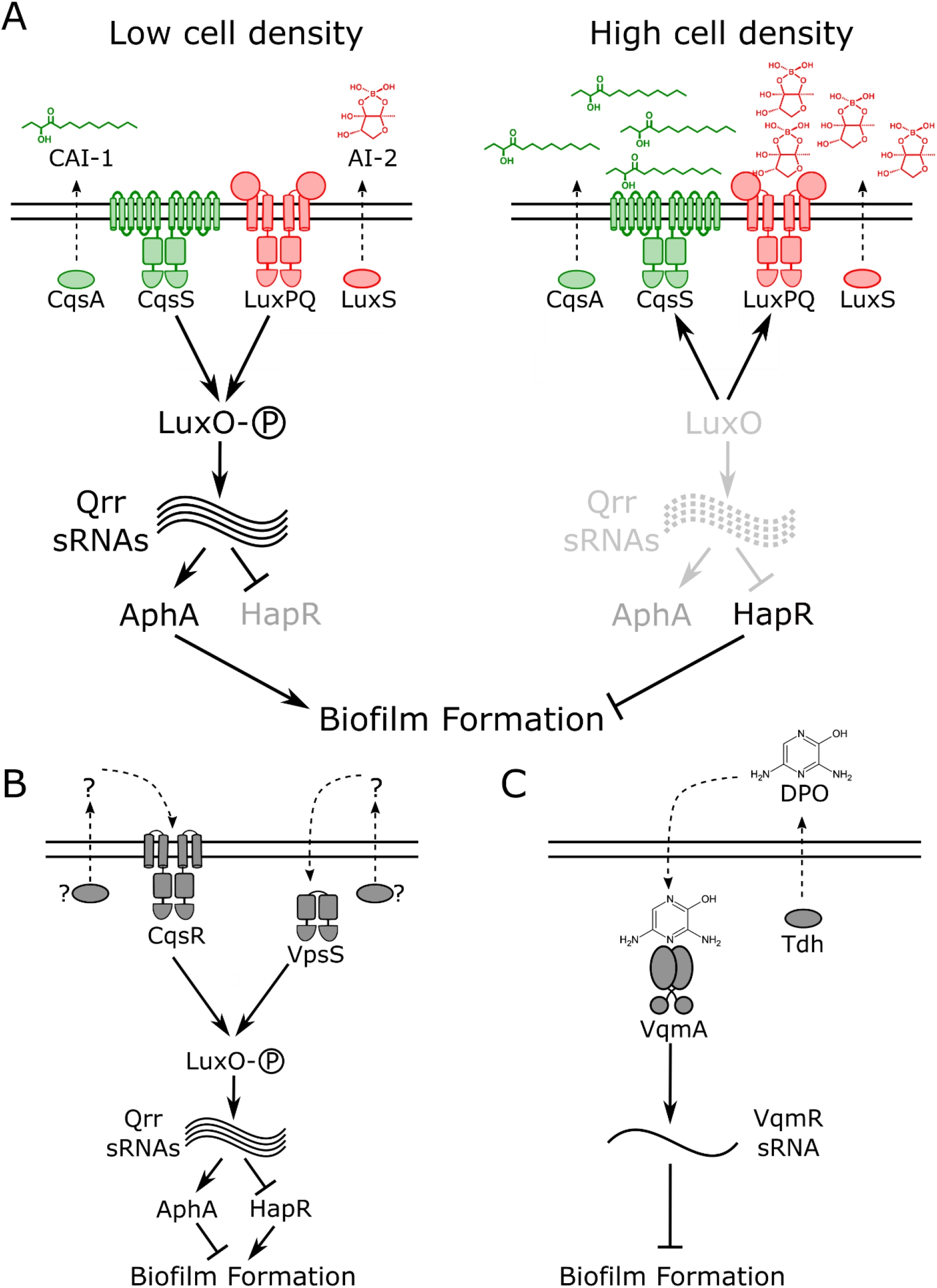
Simplified *V. cholerae* QS circuits. **(A)** Two established autoinducer-receptor pairs control QS behaviors in *V. cholerae*. One autoinducer-receptor pair consists of *cholerae* autoinducer-1 (CAI-1), synthesized by CqsA and detected by the two-component sensor-histidine kinase, CqsS. The second autoinducer-receptor pair is autoinducer-2 (AI-2), produced by LuxS and detected by LuxPQ, also a two-component sensor-histidine kinase. At LCD (left), both receptors act as kinases that promote phosphorylation of the response regulator, LuxO. LuxO∼P activates expression of genes encoding regulatory RNAs called the Qrr sRNAs. The Qrr sRNAs activate production of the LCD master regulator, AphA, and repress production of the HCD master regulator, HapR. These conditions drive biofilm formation. At HCD (right), the autoinducer-bound receptors act as phosphatases that strip phosphate from LuxO, resulting in AphA repression and HapR production, conditions that promote the free-swimming, planktonic lifestyle. (B) Two additional QS receptors, VpsS and CqsR, also funnel information into LuxO. Their cognate autoinducers and autoinducer synthases are not known. (C) A recently discovered QS pathway consisting of the autoinducer DPO, synthesized by threonine dehydrogenase (Tdh), and its partner receptor VqmA. At HCD, DPO-bound VqmA activates expression of a gene encoding a sRNA called VqmR. VqmR represses biofilm formation.

Recently, we discovered another *V. cholerae* QS pathway that functions independently of the above QS system (Fig 1C) [28–30]. In this case, the autoinducer, called DPO (3,5-dimethylpyrazin-2-ol), binds to a cytoplasmic transcriptional regulator, called VqmA. The VqmA-DPO complex activates expression of a gene encoding a regulatory RNA, called VqmR. VqmR represses genes required for biofilm formation. Thus, the DPO-VqmA-VqmR circuit also represses biofilm formation at HCD.

Here, we develop a real-time assay to measure WT *V. cholerae* biofilm formation and dispersal. The assay does not demand the use of locked QS mutants, allowing us to examine the role QS plays over the entire biofilm lifecycle. We find that the CAI-1 and AI-2 QS pathways control biofilm formation, while the DPO pathway has no effect in this assay. The AI-2 receptor LuxPQ strongly promotes biofilm formation at LCD when the ligand is absent while the CAI-1 receptor, CqsS, is incapable of driving biofilm formation at LCD. The mechanism underlying the effect stems from markedly different cell-density thresholds required for autoinducer detection by the two QS receptors, with the kin CAI-1 autoinducer threshold being achieved at much lower cell densities than that of the non-kin AI-2 autoinducer. Nonetheless, we show that both autoinducers must be present simultaneously for repression of biofilm formation to occur, suggesting that the QS system functions as a coincidence detector. Collectively, our results show that a small number of kin must be present to activate *V. cholerae* QS but the pace at which QS occurs is driven by the timing by which the inter-species AI-2 autoinducer accumulates. To our knowledge, this is the first report of unique roles for the different *V. cholerae* autoinducers, and our findings imply that detection of self fosters a different outcome than detection of other.

## Results

### A new biofilm growth and dispersal assay for WT *V. cholerae*

In *V. cholerae* biofilm studies to date, researchers have overwhelmingly employed either hyper-biofilm forming *V. cholerae* strains that are locked at LCD and incapable of QS and biofilm dispersal, or they have used fluid flow to wash autoinducers away from growing WT biofilms, in effect locking the *V. cholerae* cells at LCD [9,31–34]. While these strategies have accelerated studies of early *V. cholerae* biofilm formation and enabled identification and characterization of biofilm matrix components, QS, which is known to control the process, has not been systematically examined in a WT *V. cholerae* strain capable of naturally transitioning between LCD and HCD behavior as biofilms form and disperse. We developed simple static growth conditions that permitted WT *V. cholerae* biofilm formation and dispersal. Our strategy allows endogenously-produced autoinducers to accumulate and drive changes in QS-controlled gene expression in living, growing WT *V. cholerae* biofilms, to our knowledge, a first for the field. We used *V. cholerae*, O1 biovar El Tor strain C6706, that is known to transition between the biofilm and free-swimming states. This strain, when inoculated at LCD onto glass coverslips in minimal medium, grew into discrete biofilms, and, subsequently, biofilm dispersal occurred (Fig 2A, Movie 1). Many biofilms were simultaneously imaged over time using low-magnification brightfield microscopy. With these images, we could measure bulk biofilm biomass accumulation by performing intensity-based segmentation of the biofilms coupled with quantitation of the attenuation of light that occurred due to biofilm growth. This procedure revealed that WT biofilms grew to peak biomass at an average of ∼8-9 h after inoculation and complete dispersal occurred by ∼13 h (Fig 2B). To confirm that the imaged cell clusters were indeed biofilms, we conducted identical experiments using a Δ*vpsL* mutant strain that is incapable of producing the major polysaccharide component of the extracellular matrix required for biofilm formation [31]. No biofilm formation was detected in this mutant, validating our method (Fig 2B, Movie 1).

**Fig 2.**
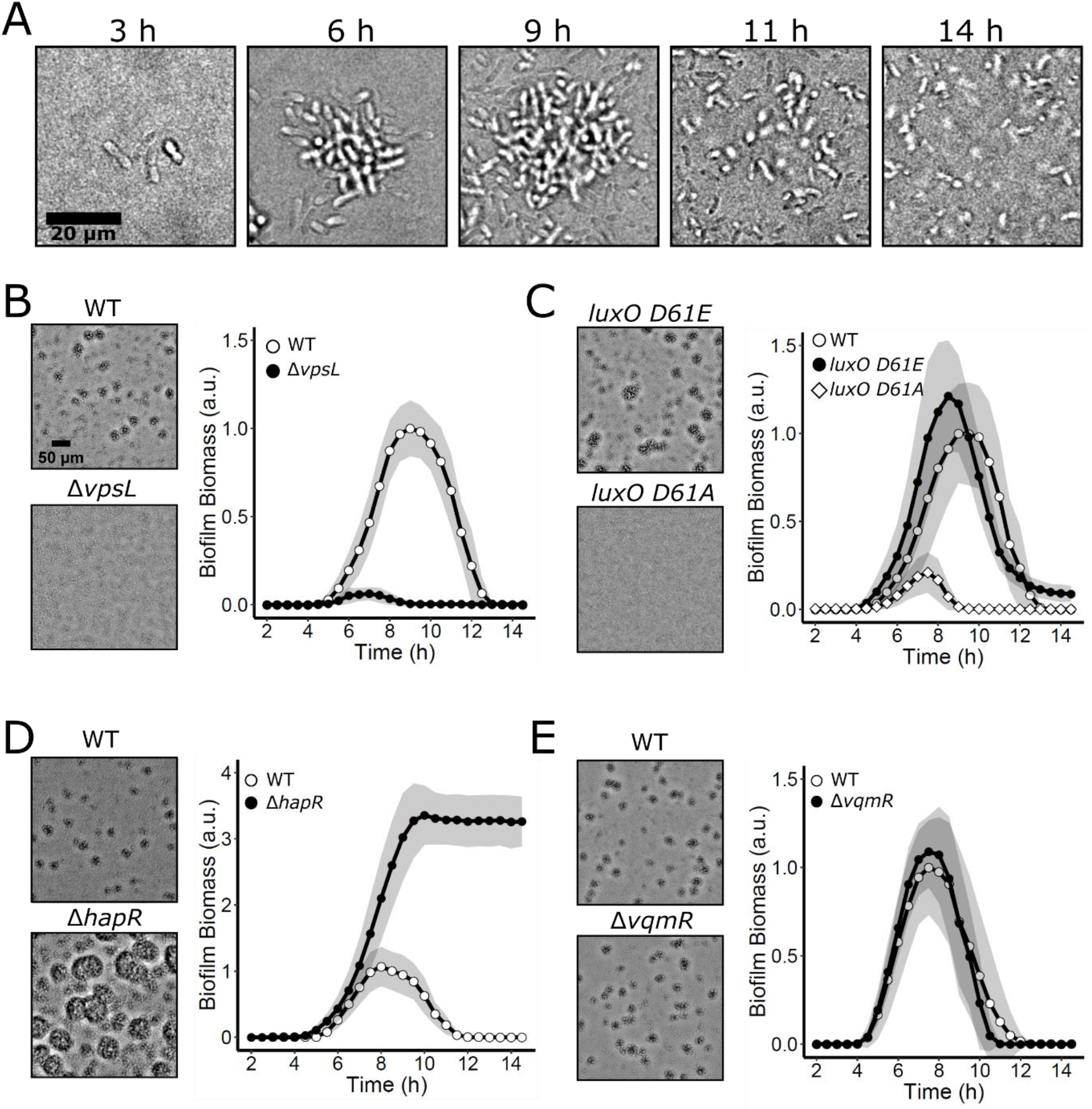
WT *V. cholerae* biofilm formation and dispersal under static growth conditions. (A) Time course of a representative WT *V. cholerae* biofilm lifecycle as imaged by bright-field microscopy using high magnification (63x objective). (B) Left panels: bright-field projections of *V. cholerae* biofilms in the indicated strains after 9 h of growth at 30°C, imaged using low-magnification (10x objective) Right panel: Quantitation of *V. cholerae* WT and Δ*vpsL* biofilm biomass over time. (C) As in B for *V. cholerae* WT and QS mutants locked in LCD (*luxO D61E*) and HCD (*luxO D61A*) modes. (D) As in B for *V. cholerae* WT and the LCD locked Δ*hapR* strain. (E) As in B for *V. cholerae* WT and the Δ*vqmR* strain. Data are represented as means normalized to the peak biofilm biomass of the WT strain in each experiment. In all cases, n=3 biological and n=3 technical replicates, ± SD (shaded).

To assess how this biofilm growth and dispersal assay compares to previous methods used for measuring QS control of *V. cholerae* biofilm formation, we analyzed the biofilm formation and dispersal phenotypes of mutant strains locked in the QS LCD and HCD modes. As mentioned, QS represses biofilm formation at HCD, and consistent with this pattern, both the LCD locked *luxO D61E* mutant carrying a LuxO∼P mimetic, and the Δ*hapR* mutant lacking the HCD master QS regulator (see Fig 1A), accumulated greater biofilm biomass than WT *V. cholerae*. Moreover, neither mutant fully dispersed (Fig 2C, D, Movie 1). Notably, the phenotype of the Δ*hapR* strain was more extreme in its preference for the biofilm state than that of the *luxO D61E* strain, consistent with the downstream position and direct function of HapR in regulation of biofilm formation. Specifically, LuxO D61E drives constitutive production of the Qrr sRNAs (Fig 1A) [35]. The Qrr sRNAs activate translation of AphA and repress translation of HapR, and they positively and negatively regulate other targets [36,37]. Thus, in the *luxO D61E* strain, unlike in the Δ*hapR* strain, some HapR is present that can activate modest biofilm dispersal, and, moreover, other Qrr-regulated targets also promote biofilm dispersal in the LuxO D61E mutant. These features of the QS circuit have been reported previously and underlie the difference in phenotypes between the two mutants [36]. A strain carrying the non-phosphorylatable *luxO D61A* allele, which is locked in the HCD QS mode failed to form appreciable biofilms (Fig 2C, Movie 1). Together, these data verify that the *V. cholerae* canonical QS system shown in Fig 1A controls biofilm formation in our assay. Below, we probe the roles of the individual QS circuits. To assess the contribution from the DPO-VqmA-VqmR pathway, we measured biofilm biomass over time in a strain lacking the VqmR regulatory RNA. At HCD, the Δ*vqmR* mutant cannot repress biofilm genes (Fig 1C). The *vqmR* mutant displayed WT biofilm formation and dispersal behaviors (Fig 2E) suggesting that the DPO-VqmA-VqmR pathway does not influence biofilm phenotypes under our assay conditions and/or when the canonical QS system is present. We do not study the DPO-VqmA-VqmR pathway further in the present work.

### AphA and HapR exhibit inverse production patterns during biofilm development, and AphA predominates in biofilms

The functioning of the canonical QS system is well established in WT *V. cholerae* cells under planktonic growth conditions: AphA is highest in abundance at LCD and its levels decline as cell density increases. Conversely, HapR is present at low levels at LCD and it accumulates with increasing cell density [26,38]. We wondered whether this inverse relationship also exists in growing WT biofilms. To examine the patterns of the two regulators, we measured the abundances of AphA and HapR during biofilm formation by building strains carrying either chromosomal *aphA-mNG* (mNeonGreen) or chromosomal *hapR-mNG* at their native loci. We also introduced a constitutive fluorescent reporter, *P_TAC_-mRuby3*, into each strain for normalization. We reasoned that the relative amounts of the AphA-mNG and mRuby3 or HapR-mNG and mRuby3 in cells could be used as a proxy for QS state. Using the above low-magnification imaging technique, coupled with confocal fluorescence microscopy and single-biofilm segmentation, we measured the fluorescence outputs from the reporters in individual biofilms over time (Fig 3). The AphA-mNG signal declined 4-fold over the lifetime of the biofilm relative to the constitutive reporter (Fig 3A, B). Conversely, following dilution of the HCD overnight culture into the biofilm assay, the HapR-mNG output decayed during early biofilm formation. Thereafter, HapR-mNG was only minimally produced until about 5-6 h of biofilm development. At that time, the HapR-mNG fluorescence signal began to increase, and it peaked immediately prior to dispersal (Fig 3C, D). Subsequently, HapR-mNG was abundantly present in cells that had become planktonic while AphA-mNG was undetectable in planktonic cells (Movie 2). The ratio of AphA-mNG:HapR-mNG throughout the time-course revealed that for the first 3.5 h of biofilm development, there was 10-17-fold more AphA than HapR (Fig 3D inset). The ratio then steadily declined, and immediately preceding dispersal, the AphA:HapR ratio was ∼1:1. We conclude that the majority of the *V. cholerae* biofilm lifetime is spent in an AphA-dominated regime. Only immediately preceding dispersal does the level of HapR increase, resulting in the transition to the planktonic lifestyle. These results suggest that AphA and HapR levels vary inversely in response to changes in cell density, and that relationship is maintained in both biofilm and planktonic cells. Thus, the core behavior of the QS system is conserved in both growth modes.

**Fig 3.**
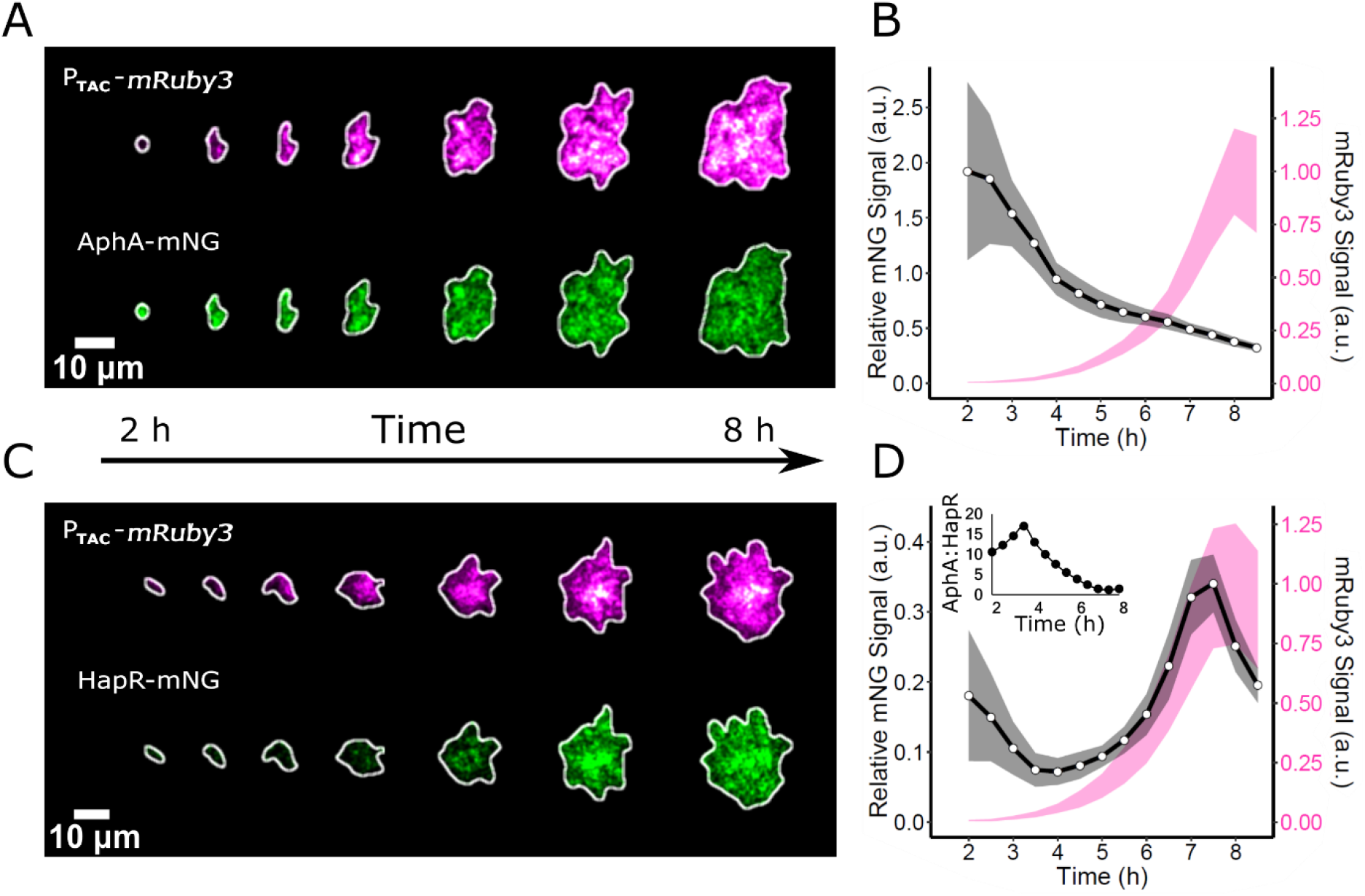
AphA and HapR abundances vary inversely during biofilm formation. (A) Representative image series showing the formation of an individual biofilm harboring the constitutive reporter P_TAC_-*mRuby3* and AphA-mNG (mNeonGreen). (B) Quantitation of the AphA-mNG fluorescence (black line) relative to the control mRuby3 fluorescence (magenta line) over the course of biofilm development. n=24 biofilms from 3 biological replicates. (C and D) As in A and B, respectively, for HapR-mNG. Inset in (D) represents the AphA-mNG:HapR-mNG ratio over time. Shading in B and D represents SD.

### AI-2 represses WT *V. cholerae* biofilm formation

We wondered how autoinducers influence *V. cholerae* biofilm formation and dispersal. As mentioned, two QS receptors, VpsS and CqsR, have recently been discovered that feed information into the canonical QS pathway but their cognate autoinducers and autoinducer synthases are not identified (Fig 1B) [17]. For that reason, we cannot control autoinducer production for these two circuits nor can we quantify their inputs into QS-driven biofilm behavior. To avoid confounding issues arising from signaling by two unidentified autoinducers, in some experiments, we deleted the *vpsS* and *cqsR* genes so that inputs from the two unknown cues were eliminated, allowing us to quantitatively assess the activities of CAI-1 and AI-2. In every experiment, we specify whether the *vpsS* and *cqsR* genes are present or not.

To probe the individual roles of CAI-1 and AI-2 in repression of biofilm formation and driving biofilm dispersal, we built reporter strains that exclusively respond to only one of these two autoinducers. Each reporter strain possesses a single QS receptor, but it lacks the corresponding autoinducer synthase. Thus, only exogenously supplied autoinducer can activate QS, and only via the single remaining receptor. To our surprise, addition of synthetic CAI-1 at a saturating concentration of 5 μM (EC_50_ = 32 nM, [39]), at the initiation of biofilm formation had no effect on biofilm development or dispersal in the CAI-1 reporter strain (Δ*vpsS*, Δ*cqsR*, Δ*luxQ*, Δ*cqsA*) as results were identical to when solvent was added (S1 Fig). In contrast, administration of 5 μM of a structurally unrelated CqsS agonist (EC_50_ = 9 nM, [39]), 1-ethyl-*N*-{[4-(propan-2-yl)phenyl]methyl}-1*H*-tetrazol-5-amine, (that we call CAI-1* for simplicity), markedly reduced biofilm formation (S1A Fig). We confirmed that our synthetic CAI-1 is fully active in this reporter strain by monitoring bioluminescence emission from a chromosomally integrated luciferase (*lux*) operon, a routinely-used heterologous readout for HapR-controlled QS activity in *V. cholerae* [27]. When grown in shaken, planktonic conditions, both CAI-1 and CAI-1* induced an ∼1000-fold increase in light production by the CAI-1 reporter strain, suggesting that synthetic CAI-1 is only inactive, possibly due to micelle formation, under biofilm growth conditions (S1B, C Fig). For this reason, we use CAI-1* in place of CAI-1 for the remainder of this work. Addition of saturating AI-2 (5 μM; EC_50_ = 21 nM as measured in *Vibrio harveyi* [16]) to the corresponding AI-2 reporter strain (Δ*vpsS*, Δ*cqsR*, Δ*cqsS*, Δ*luxS*) dramatically reduced biofilm formation and, moreover, activated the *lux* reporter ∼1000-fold, showing that AI-2 is active in both assays (S2A, B Fig, respectively).

We next explored how exogenous provision of CAI-1* or AI-2 influences the WT *V. cholerae* biofilm program, in the case in which all four QS receptors are present and all of the autoinducers are also endogenously produced and accumulate naturally over time. The architecture of the *V. cholerae* QS system is arranged such that all four autoinducers feed information into the same signal integrator, LuxO, and as such, the expectation is that administration of additional CAI-1* or AI-2 should prevent biofilm formation and/or promote dispersal (Fig 1A). To the contrary, we found that the addition of 5 μM CAI-1* to *V. cholerae* cells that naturally produce CAI-1 and AI-2 had little effect on WT biofilm biomass accumulation or dispersal (Fig 4A, B). However, addition of 5 μM AI-2 repressed biofilm formation and promoted premature biofilm dispersal (Fig 4A, B). We obtained identical results when the *vpsS* and *cqsR* genes were present and when they had been deleted, showing that input from these two circuits is negligible under these conditions (S3 Fig). To confirm that AI-2 caused its effect via the *V. cholerae* QS system, we assayed whether AI-2 could repress biofilm formation in the *V. cholerae luxO D61E* strain that is locked in the LCD QS mode and is impervious to autoinducers [35]. AI-2 had no effect on biofilm formation or dispersal in this strain (Fig 4C). Therefore, AI-2 requires a functional QS system to drive changes in *V. cholerae* biofilm behavior. These results suggest that exogenously-supplied AI-2 but not CAI-1* should foster premature induction of HapR, the downstream master regulator of the QS HCD state. Indeed, saturating AI-2 caused HapR-mNG production to increase after only 3 h of biofilm growth, and by 6 h, HapR-mNG levels were 8-fold higher than in untreated biofilms and 3-fold higher than in CAI-1* treated biofilms (Fig 4D, E). To our knowledge, these findings represent the first case in which AI-2/LuxPQ activity has a stronger effect than CAI-1/CqsS activity on *V. cholerae* QS behavior.

**Fig 4.**
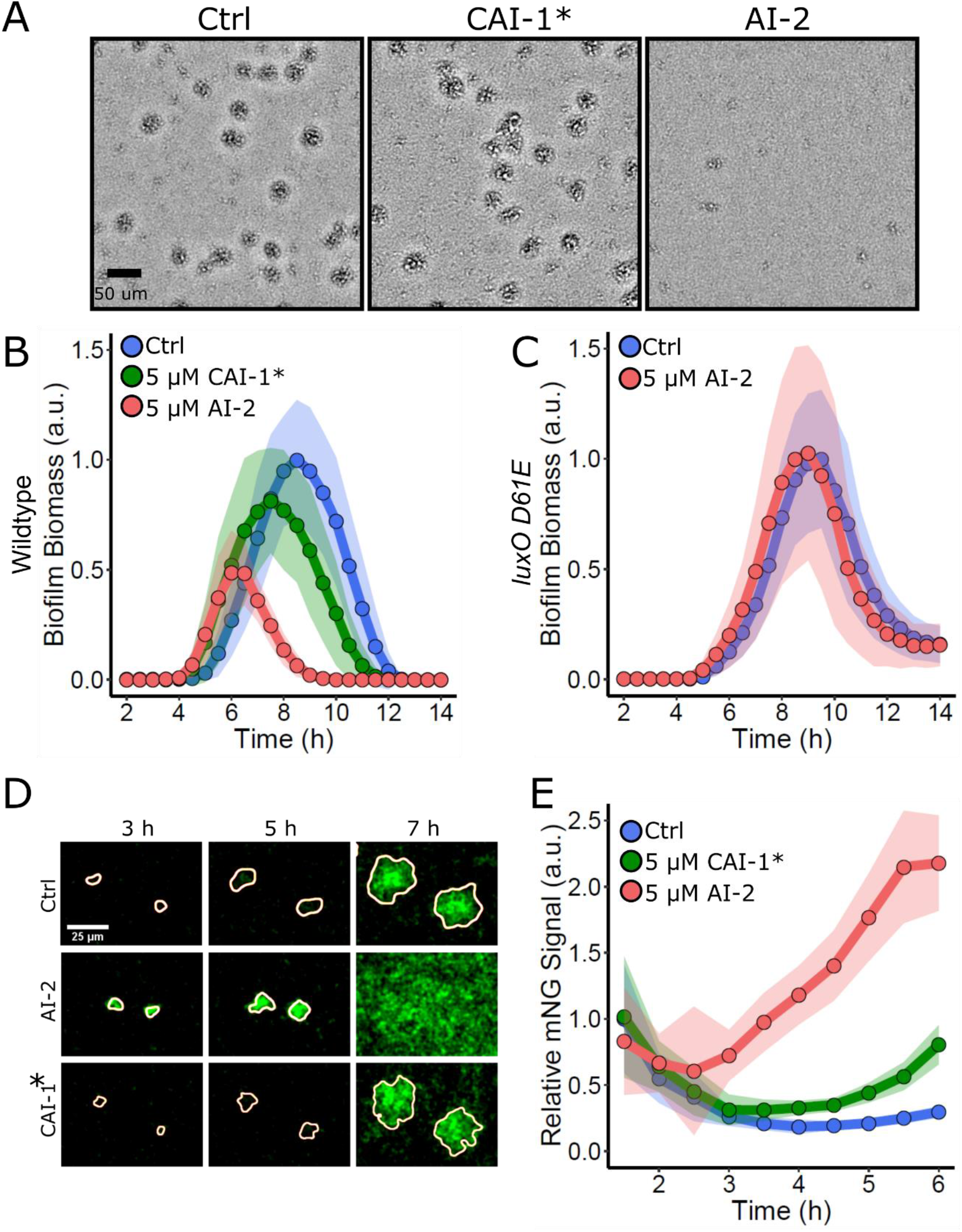
Exogenous AI-2 but not CAI-1* represses WT *V. cholerae* biofilm formation. (A) Representative projections of WT *V. cholerae* treated with 0.25% DMSO (Ctrl), 5 μM CAI-1*, or 5 μM AI-2 after 9 h of biofilm growth at 30°C. (B) Quantitation of biofilm biomass for WT *V. cholerae* treated with 0.25% DMSO (Ctrl), 5 μM CAI-1*, or 5 μM AI-2 over time. Data are represented as means normalized to the peak biofilm biomass of the DMSO control. In all cases, n=3 biological and n=3 technical replicates, ± SD (shaded). (C) As in B for the *V. cholerae luxO D61E* strain treated with 0.25% DMSO (Ctrl) or 5 μM AI-2. (D) Representative images of WT *V. cholerae* producing HapR-mNG after treatment as in B. (E) Quantitation of HapR-mNG signal relative to the control, P_TAC_-*mRuby3* signal over the course of biofilm development following treatment as in B. n=24 biofilms from 3 biological replicates. Data are normalized to the initial intensity of the sample to which DMSO was added.

The differences in strengths of the CAI-1 and AI-2 autoinducers on biofilm repression were unexpected. We wondered whether the dominance of the AI-2 signal was specific to the biofilm mode of growth or if other traits, deployed in conventional, shaken, planktonic growth conditions would likewise be differentially controlled. To test this possibility, we monitored expression of the chromosomally-integrated *lux* operon following administration of autoinducers. In contrast to biofilm formation, which is repressed by HapR at HCD, autoinducer accumulation drives HapR to activate *lux* gene expression at HCD. Specifically, in bioluminescence assays, light output is high immediately following dilution of an HCD overnight planktonic culture. Thereafter, light production declines precipitously because the autoinducers have been diluted to below their levels of detection. As the cells grow, endogenously-produced autoinducers accumulate, and light production again commences. Thus, a “U” shaped curve is a hallmark QS-activated gene expression pattern (S4A Fig). To examine the effect of each autoinducer on *lux* activation, we administered 5 μM of either CAI-1* or AI-2 to WT *V. cholerae* carrying the *lux* reporter. The results mirror those shown for biofilm formation in Fig 4B except the result is activation not repression of behavior. Here, at LCD, addition of AI-2 but not CAI-1* stimulated a 10-fold enhancement in light production irrespective of whether the *vpsS* and *cqsS* genes are present or not (S4B, C Fig). Together, our results exploring QS repression of biofilm formation and activation of light production show that WT *V. cholerae* is impervious to exogenously supplied CAI-1* but is exquisitely sensitive to exogenously supplied AI-2.

### CqsS is activated at extremely low cell densities

Given that LuxPQ and CqsS relay information to the same response regulator, LuxO, it was not obvious how exogenous AI-2 could so dominate the WT QS phenotypes. First, regarding biofilms: one possibility is that LuxPQ kinase activity is required for biofilm formation at LCD while CqsS kinase activity is dispensable. If so, exogenous AI-2, but not CAI-1* would drive repression of biofilm formation and activation of biofilm dispersal. To test this idea, we measured biofilm formation and dispersal in strains possessing only a single autoinducer synthase-receptor pair, either LuxS/AI-2 and LuxPQ (designated AI-2^S+R+^) or CqsA/CAI-1 and CqsS (designated CAI-1^S+R+^) and compared them to the strain containing both synthase-receptor pairs (designated CAI-1^S+R+^, AI-2^S+R+^) In in all cases, the strains lacked the VpsS and CqsR receptors. (See the schematic in Fig 5A for the depiction of the strains.) Importantly, in these experiments, we did not supply exogenous autoinducers. The AI-2^S+R+^ strain accumulated biofilm biomass and dispersed identically to the CAI-1^S+R+^, AI-2^S+R+^ strain (Fig 5A, middle panel). By contrast, the CAI-1^S+R+^ strain was defective in the ability to form biofilms and it dispersed prematurely (Fig 5A, middle panel). This experiment shows that the LuxPQ kinase can drive *V. cholerae* biofilm formation at LCD while the CqsS kinase cannot. To determine if this relationship is unique to biofilm growth, or if it also applies to planktonic behaviors, we measured the ability of the same strains to activate *lux* expression. The AI-2^S+R+^ strain showed the WT (i.e., CAI-1^S+R+^, AI-2^S+R+^) pattern for light production (Fig 5A, right panel). By contrast, at LCD, the CAI-1^S+R+^ strain produced 100-fold more light than the AI-2^S+R+^ and CAI-1^S+R+^, AI-2^S+R+^ strains, resulting in a pattern of light production nearly indistinguishable from a strain lacking the four QS receptors. The mutant that has no QS receptors lacks all QS kinase inputs and therefore produces maximal constitutive bioluminescence (Fig 5A, right panel, depicted in black). Together, these results show that LuxPQ establishes the LCD QS mode while the CqsS receptor does not do so in biofilms or in planktonic cells.

**Fig 5.**
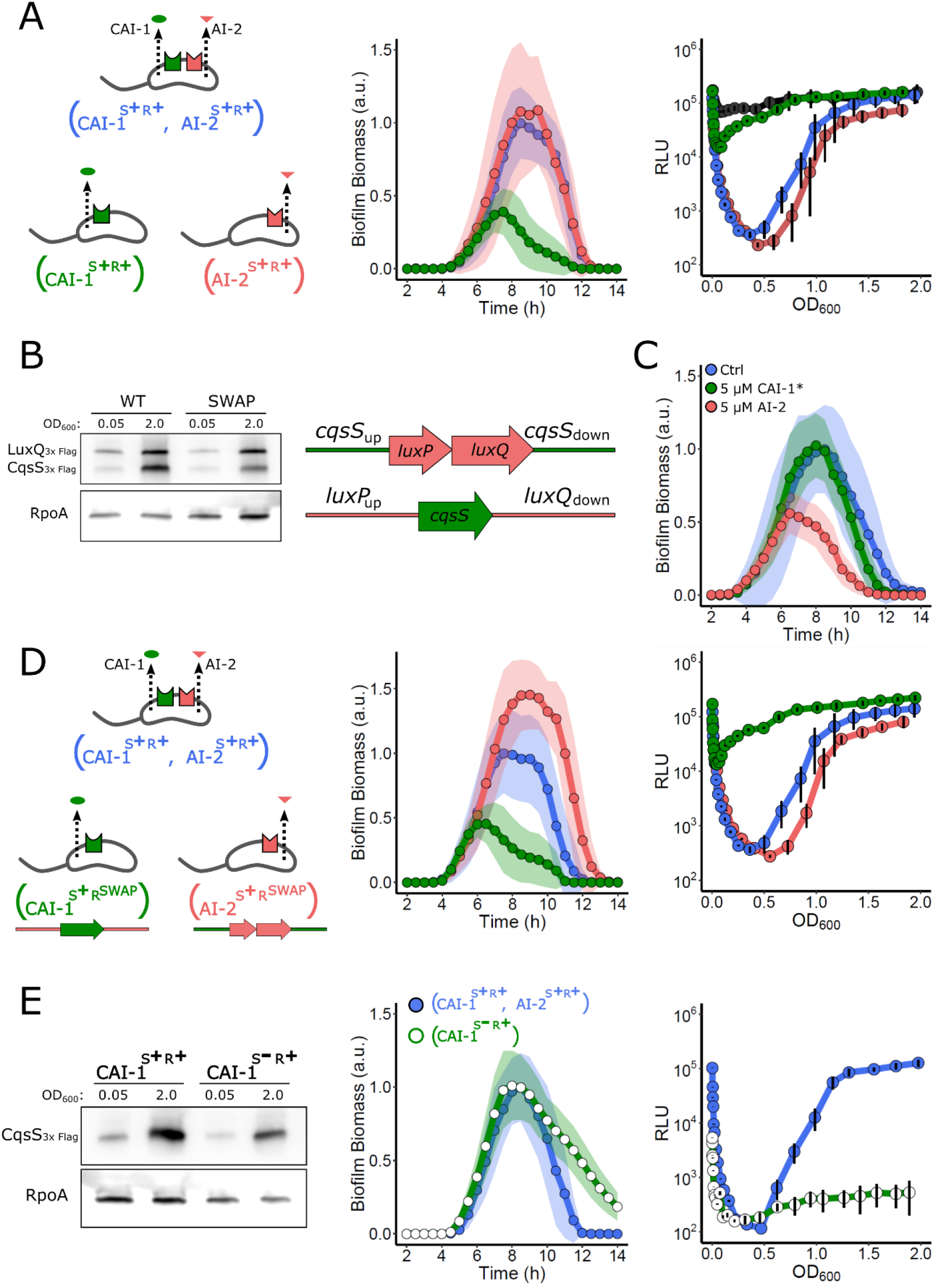
LuxPQ but not CqsS drives LCD QS behaviors. (A) Left panel: Schematic representing *V. cholerae* strains containing both QS circuits or that produce and detect a single autoinducer. Middle panel: Quantitation of biofilm biomass over time for the strain possessing both QS systems (AI-2^S+R+^, CqsS^S+R+^; blue), only the AI-2 QS system (AI-2^S+R+^ red), and only the CAI-1 QS system (CqsS^S+R+^; green). Right panel: The corresponding *lux* patterns for the strains in the middle panel. The additional black curve shows the result for the *V. cholerae* strain lacking all four QS receptors (Δ*vpsS*, Δ*cqsR*, Δ*luxQ*, Δ*cqsS*). (B) Left panel: Representative western blot for a strain containing 3X-FLAG tagged CqsS and LuxPQ produced from their native loci (WT) and for a strain in which their genomic positions had been exchanged (SWAP). RpoA was used as the loading control. Right panel: Schematic showing exchange of the *cqsS* and *luxPQ* genomic locations. (C) Quantitation of biofilm biomass for the strain with the exchanged LuxPQ and CqsS alleles (CqsS^S+RSWAP^, AI-2^S+RSWAP^) treated with 0.25% DMSO (Ctrl), 5 μM CAI-1*, or 5 μM AI-2 over time. (D) Left panel: Schematic representing *V. cholerae* strains that contain both QS circuits or that produce and detect a single autoinducer in which the receptor genes are expressed from the exchanged loci. Middle panel: Quantification of biofilm biomass over time for a *V. cholerae* strain possessing both QS systems (CAI-1^S+R+^, AI-2^S+R+^; blue), the AI-2 system only, with *luxPQ* expressed from the *cqsS* locus (AI-2^S+RSWAP^; red), and the CAI-1 system only, with *cqsS* expressed from the *luxPQ* locus (CAI-1^S+RSWAP^; green). Right panel: The corresponding *lux* patterns for the strains in the middle panel. (E) Left panel: Representative western blot showing CqsS-3XFLAG levels in the *V. cholerae* CAI-1^S+R+^ and CAI-1^S-R+^ strains. Middle panel: Quantitation of biofilm biomass over time for the *V. cholerae* CAI-1^S+R+^, AI-2^S+R+^ (blue circles) and CAI-1 ^S-R+^ (green circles) strains. Right panel: The corresponding *lux* patterns for the strains in the middle panel. In all biofilm measurements, data are represented as means normalized to the peak biofilm biomass of the CAI-1^S+R+^, AI-2^S+R+^ strain and n=3 biological and n=3 technical replicates, ± SD (shaded). In all *lux* experiments, relative light units (RLU) are defined as light production (a.u.) divided by OD_600_ and n=3 biological replicates, and error bars represent SD.

One mechanism that could underlie the, respectively, strong and weak effects of LuxPQ and CqsS in control of LCD QS behaviors is that *cqsS* is not sufficiently expressed at LCD, effectively making CqsS absent and therefore unable to promote the LCD QS state. If, by contrast, LuxPQ is present at LCD, its kinase could be exclusively responsible for promoting LCD QS behaviors. Western blot analysis of a strain containing 3X-FLAG tagged LuxQ and 3X-FLAG tagged CqsS produced from their native loci revealed that LuxQ was roughly twice as abundant as CqsS at LCD, while CqsS was in excess of LuxQ at HCD (Fig 5B, left panel). We next exchanged the genomic positions of *cqsS-3X-FLAG* and *luxPQ-3X-FLAG*, placing each receptor gene under the other’s promoter. (Fig 5B, schematic). In this case, LuxQ and CqsS were present at approximately equal levels at LCD and LuxQ was in excess of CqsS at HCD (Fig 5B, left panel). Provision of exogenous AI-2 or CAI-1* to the strain containing the exchanged alleles (CAI-1^S+RSWAP^, AI-2^S+RSWAP^) revealed that AI-2 remained the dominant autoinducer in LCD repression of biofilm formation (Fig 5C). Moreover, in strains carrying a single synthase-receptor pair in which the genomic locations of the receptors had been exchanged (designated CAI-1^S+RSWAP^ and AI-2^S+RSWAP^; see schematic in Fig 5D), little biofilm formation occurred when *cqsS* was expressed from the *luxPQ* locus, while biofilm biomass accumulated in excess of that in WT *V. cholerae* when *luxPQ* was expressed from the *cqsS* locus (Fig 5D, middle panel). Consistent with this finding, in the luciferase assay, the strains containing the singly exchanged receptors behaved the same as when the respective receptor gene was expressed from its native site (Fig 5D, right panel). These results show that the WT relative abundances of the QS receptors cannot explain the difference between the CqsS and LuxPQ kinase activities, and in turn, their influence on QS at LCD.

We considered two other possibilities to explain the variation in QS receptor kinase activity at LCD. First, either CqsS is an intrinsically poor kinase when unliganded, so it cannot drive the LCD state, or second, CqsS binds to the CAI-1 autoinducer and switches from kinase to phosphatase mode at cell densities much lower than those traditionally considered to be LCD, so its influence over the LCD QS state is rapidly abolished as the cells grow. To distinguish between these two possibilities, we deleted the CAI-1 autoinducer synthase gene, *cqsA*, from the CAI-1^S+R+^ strain, generating the CAI-1^S-R+^ strain, and we examined the ability of this strain to establish the LCD behavior. Importantly, the amount of CqsS present at LCD, as measured by western blotting, was identical in the CAI-1^S-R+^ strain and that of the CAI-1^S+R+^ parent strain that contains *cqsA* (Fig 5E, left panel). The CAI-1^S-R+^ strain was capable of driving WT levels of biofilm formation and, moreover, exhibited a delay in dispersal (Fig 5E, middle panel). Furthermore, the CAI-1^S-R+^ strain failed to activate light production in the planktonic *lux* assay irrespective of cell density (Fig 5E, right panel). These data demonstrate that, when the CAI-1 autoinducer is absent, the CqsS kinase is indeed sufficiently potent to drive the LCD QS program both on surfaces and in planktonic conditions. Thus, in strains that possess both CqsA and CqsS, at LCD, there must be enough CAI-1 autoinducer present to inhibit CqsS kinase-driven biofilm formation and prevent *lux* expression.

### The *V. cholerae* QS system is a coincidence detector

Based on the above results, we suggest that, at very low cell densities, sufficient CAI-1 is present to bind the CqsS receptor and convert it from kinase to phosphatase mode. By contrast, because the critical concentration of AI-2 required to transform LuxPQ from a kinase to a phosphatase is achieved only at higher cell densities, LuxPQ remains a kinase enabling biofilms to form and begin to mature, and in the case of luciferase, *lux* is not activated. If so, during this time window, the activities of the two receptors oppose one another. We know that kinase activity is critical for establishing the LCD QS program, and since biofilms form, and light production is off at LCD, it suggests that LuxPQ kinase overrides CqsS phosphatase. Following this same logic, we hypothesize that, if kinase activity must dominate for LCD behaviors to be undertaken, it should not matter which receptor is the kinase and which receptor is the phosphatase. To test this supposition, we measured light output in a *V. cholerae* strain possessing both QS receptors, but lacking the AI-2 synthase, LuxS (CAI-1^S+R+^, AI-2^S-R+^). In this case, CqsS switches from kinase to phosphatase upon CAI-1 binding and LuxPQ is a constitutive kinase. The CAI-1^S+R+^, AI-2^S-R+^ strain produced ∼1000-fold less light than the CAI-1^S+R+^, AI-2^S+R+^ strain that contains both autoinducer-receptor pairs (Fig 6A). We performed the reciprocal experiment using a strain lacking the CAI-1 synthase, CqsA (CAI-1^S-R+^, AI-2^S+R+^). In this case, CqsS is the constitutive kinase and LuxPQ transitions from kinase to phosphatase upon AI-2 binding. This strain also exhibited 1000-fold reduced light production at LCD relative to the CAI-1^S+R+^, AI-2^S+R+^ strain (Fig 6A). Together, these results show that, kinase activity, irrespective of which receptor provides it, overrides phosphatase activity at LCD. Moreover, it means that both autoinducers must be present simultaneously for a robust and timely transition from LCD to HCD to occur. Thus, the *V. cholerae* QS system functions as a coincidence detector for the two autoinducer inputs.

**Fig 6.**
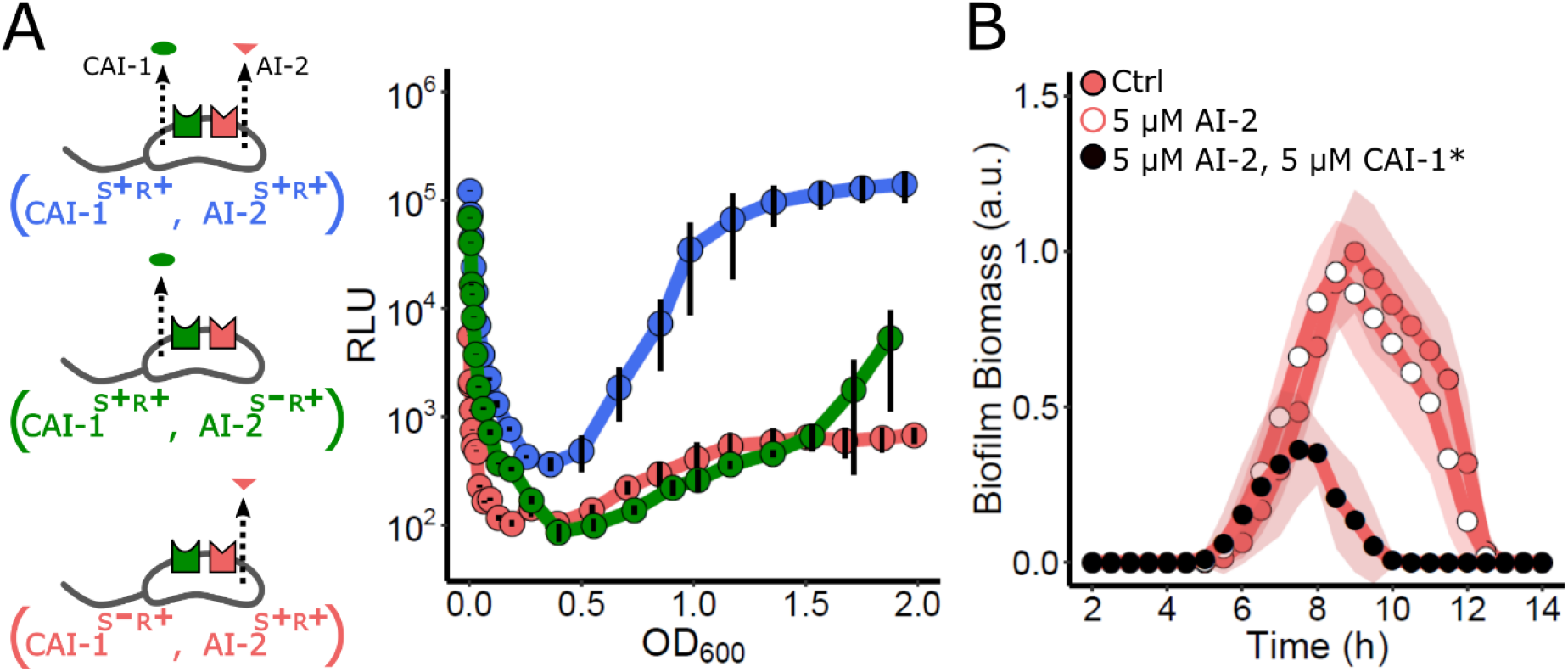
The *V. cholerae* QS circuit is a coincidence detector. (A) Left panel: Schematic for strains used in the right panel, which shows the *lux* expression patterns. The strains are: CqsS^S+R+^, AI-2^S+R+^ (blue), CqsS^S+R+^, AI-2^S-R+^ (green), and CqsS^S-R+^, AI-2^S+R+^ (red). Relative light units (RLU) are defined as light production (a.u.) divided by OD_600_. n=3 biological replicates and error bars represent SD. (B) Quantitation of biofilm biomass for the CAI-1^S-R+^, AI-2^S+R+^ strain to which DMSO solvent (red circles, Ctrl), 5 μM AI-2 (white circles), or 5 μM AI-2 and 5 μM CAI-1* (black circles) was added. Data are represented as means normalized to the peak biofilm biomass of the control and n=3 biological and n=3 technical replicates, ± SD (shaded).

With a coincidence detection model in mind, we predicted that the addition of exogenous AI-2 should have no effect on biofilm formation and dispersal in a strain possessing both receptors but lacking the CAI-1 synthase CqsA (CAI-1^S-R+^, AI-2^S+R+^). In this setup, LuxPQ would function as a phosphatase upon binding to AI-2 and the lack of the *cqsA* gene would ensure that CqsS remains a kinase at all cell densities. Thus, CqsS kinase should override AI-2-bound LuxPQ phosphatase. Indeed, Fig 6B shows that AI-2 has no effect on biofilm formation/dispersal in this strain. By contrast, simultaneous administration of CAI-1* and AI-2 to the CAI-1^S-R+^, AI-2^S+R+^ strain satisfies the coincidence detector requirement, converts both CqsS and LuxPQ to phosphatase mode, and causes biofilm repression (Fig. 6B). We conclude that while the *V. cholerae* QS system is a coincidence detector, the consequence of the exceedingly low cell density activation of CqsS by endogenously-produced CAI-1 makes it so that endogenous accumulation or exogenous sources of AI-2 satisfy the coincidence detector leading to HCD behaviors.

### CAI-1 activation of CqsS occurs via quorum sensing, not self sensing

Recently, several bacterial QS circuits have been shown to be capable of self sensing, in which an individual cell releases and detects its own autoinducer molecule by an autocrine-like mechanism (Fig 7A) [40,41]. For self sensing to occur, the cells must harbor sufficient levels of the receptor to capture/bind the released molecule prior to it diffusing away [42]. We considered the possibility that CAI-1 could be sensed by the same cell that secretes it, potentially explaining how the CAI-1/CqsS arm of the QS system becomes activated at such low cell densities relative to the AI-2/LuxPQ circuit. On the other hand, we expected that the AI-2/LuxPQ circuit must display QS behavior, rather than self sensing, explaining why, relative to the CAI-1/CqsS circuit, the AI-2/LuxPQ arm does not engage until much higher cell densities. To explore these ideas, we examined self versus non-self sensing in each circuit by co-culturing a “secrete-and-sense” strain (containing a single autoinducer synthase-receptor pair) with a “sense-only” strain (containing only that receptor) (Fig 7A). The rationale is that, if self sensing occurs, in co-culture, the autoinducer made by the secrete-and-sense strain would trigger its HCD mode, while the sense-only strain would remain in LCD mode (Fig 7B, top). By contrast, if released autoinducer is shared between the two strains, then both the secrete-and-sense and the sense-only strains would proceed through the LCD to HCD QS program simultaneously (Fig 7B, bottom).

**Fig 7.**
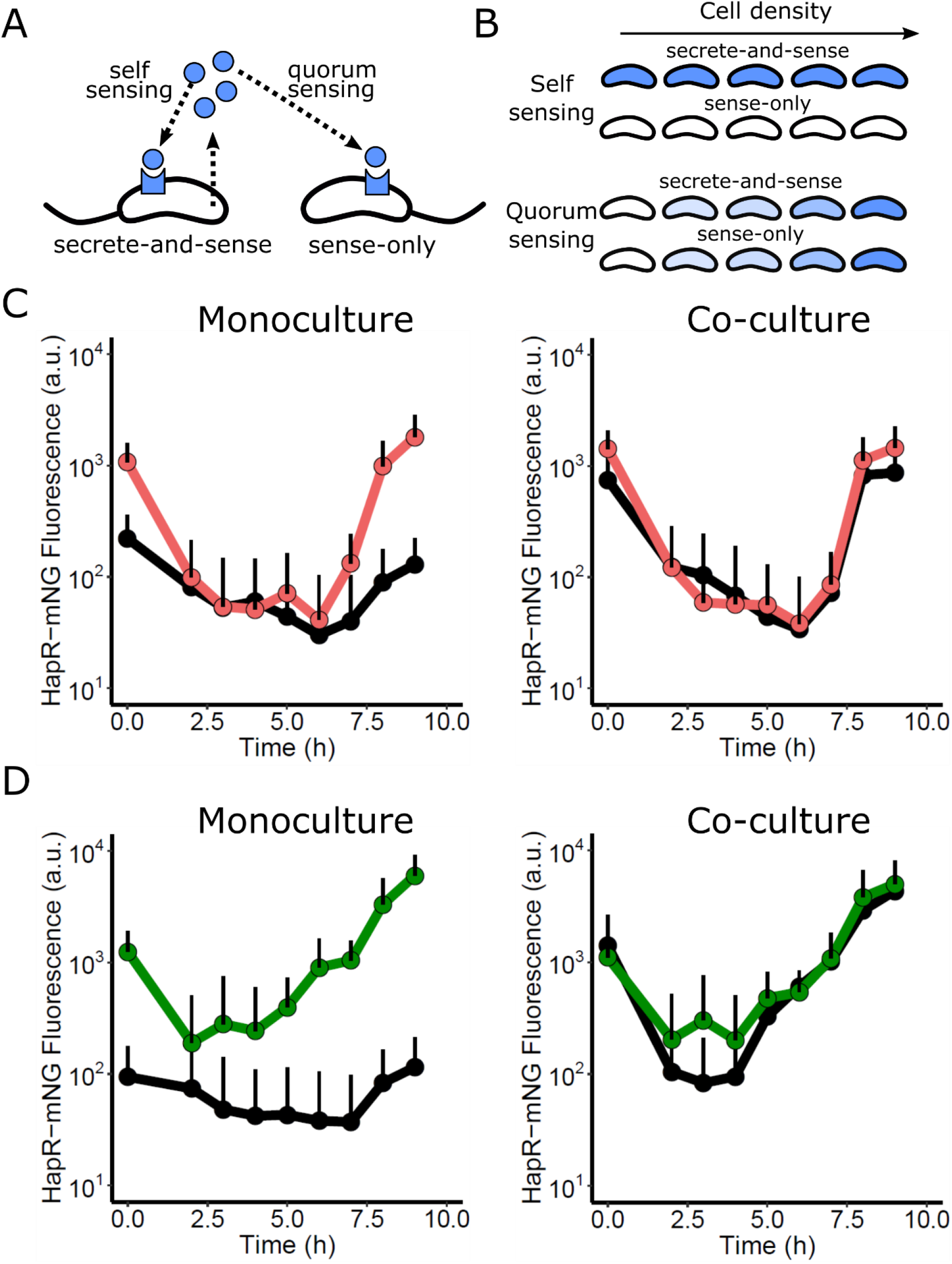
The CqsS/CAI-1 system is a QS system not a self-sensing system. (A) Schematic showing self sensing and QS. See text for details. (B) Top panel: Predicted gene expression patterns in a self-sensing system. Bottom panel: Predicted gene expression patterns in a QS system. (C) Left panel: Average individual cell HapR-mNG fluorescence for the *V. cholerae* AI-2^S+R+^ (red) and the AI-2^S-R+^ (black) strains grown in monoculture. Right panel: The same strains grown in co-culture. (D) Left panel: Average individual cell HapR-mNG fluorescence for the *V. cholerae* CAI-1^S+R+^ (green) and the CAI-1^S-R+^ (black) strains grown in monoculture. Right panel: The same strains grown in co-culture. Error bars represent SD of individual cell measurements at each timepoint.

We first examined self sensing in the AI-2/LuxPQ circuit. To characterize individual cell responses following co-culture, we used flow cytometry analyses to measure HapR-mNG fluorescence as a readout of HCD in the secrete-and-sense and sense-only strains. We differentiated between the strains by introducing a constitutive mRuby3 fluorescence reporter into one of the strains. As controls, we measured production of HapR-mNG in the AI-2/LuxPQ secrete-and-sense strain (AI-2^S+R+^), and in the sense-only strain (AI-2^S-R+^) grown in monoculture. (Fig 7C). In these experiments, we diluted the cells to the very low cell density of OD_600_ = 5 × 10^-6^, or ∼2,000 cells/mL. For reference, typical *V. cholerae* QS assays are initiated at OD_600_ = 5 × 10^-4^, or ∼200,000 cells/mL. Upon dilution, the secrete-and-sense AI-2^S+R+^ strain repressed HapR-mNG production ∼10-fold, and importantly, to the same level as the sense-only, AI-2^S-R+^ strain. In the AI-2^S+R+^ secrete-and-sense strain, HapR-mNG production remained low for many growth cycles and began to increase only after 7 h of growth, at an ∼OD_600_ of 0.1 (Fig 7C, left). In co-culture, clear QS behavior occurred: HapR-mNG fluorescence in the sense-only AI-2^S-R+^ strain matched that of the secrete-and-sense AI-2^S+R+^ strain (Fig 7C, right). These results indicate that released AI-2 is detected equally by all cells irrespective of whether or not they can produce the autoinducer.

We next performed analogous experiments to test self sensing versus QS by the CAI-1/CqsS circuit. We first analyzed the secrete-and-sense (CAI-1^S+R+^) and sense-only (CAI-1^S-R+^) strains grown in monoculture (Fig 7D, left). In this case, the secrete-and-sense CAI-1^S+R+^ strain repressed HapR-mNG 5-fold by 2 h post-dilution. Thereafter, HapR-mNG fluorescence rapidly increased. Notably, however, the secrete-and-sense CAI-1^S+R+^ strain did not repress HapR-mNG to the level of that by the sense-only CAI-1^S-R+^ strain grown alone, indicating that an even greater dilution is required to completely convert CqsS to the kinase mode. In co-culture, HapR-mNG fluorescence followed the same trajectory for the sense-only CAI-1^S-R+^ and the secrete-and-sense CAI-1^S+R+^ strains (Fig 7D, right) indicating that QS, not self sensing is the major driver of HapR-mNG induction in this circuit despite its early activation. From these results, we can conclude that self sensing does not occur in either *V. cholerae* QS circuit, however, the two circuits are activated by their cognate autoinducers at radically different cell densities, with the CAI-1/CqsS arm being activated at much lower cell densities than the AI-2/LuxPQ circuit.

## Discussion

In this study, we present a real-time assay for WT *V. cholerae* biofilm growth and dispersal. This approach enables analysis of WT *V. cholerae* that naturally transitions from LCD to HCD, and therefore progresses through the entire QS cycle. LCD locked QS *V. cholerae* mutants were analyzed in earlier iterations of biofilm assays because their constitutive hyper-biofilm-forming phenotypes enabled imaging of biofilms as they formed. The locked LCD QS mutants were especially instructive, yielding the major matrix components and their roles, cell packing patterns, and the contributions of mechanics to biofilm morphology [4,32,33]. However, the locked LCD mutants precluded assessment of QS control over the biofilm program, and, furthermore, the locked LCD mutants used in the earlier studies do not disperse from biofilms so the second part of the lifecycle – the transition from the biofilm to the planktonic phase – could not be accessed. Our new assay permits the study of the full biofilm program from initiation to dispersal and moreover, mutants that are defective in particular QS components can be studied, individual cell and bulk measurements can be made, autoinducers and analogs can be supplied exogenously, and reporter genes can be monitored individually or in combination. Furthermore, this assay is easily adapted to high-throughput microscopy approaches, as it is performed in 96-well plates and does not require the complexities of microfluidics to deliver flow. Going forward, our intention is to use the assay with a focus on the understudied dispersal process: identifying the dispersal cue(s), the genes that orchestrate dispersal, and the molecular mechanisms that enable cells to escape from matrix-covered sessile communities.

Using this new assay, we first confirmed that WT *V. cholerae* forms biofilms at LCD and disperses from them at HCD. We quantitatively imaged the master regulators to assess QS states in developing and dispersing biofilms. We found that the AphA-driven LCD regime spans nearly the entirety of the *V. cholerae* biofilm lifecycle. Control is passed to HapR, the HCD regulator, only immediately preceding biofilm dispersal. Investigation of the individual and collective roles of the kin (CAI-1) and non-kin (AI-2) receptors showed that they function as a coincidence detector: both autoinducers must be present simultaneously for repression of biofilms and launching of dispersal to occur.

Most surprising was our finding that, in growing biofilms, a marked asymmetry exists in QS signaling. Endogenously produced CAI-1 accumulates rapidly, activating the CqsS phosphatase early in biofilm development, whereas AI-2 does not accumulate to the threshold required to transition LuxPQ from a kinase to a phosphatase until biofilms are significantly more mature (Fig 8A). Thus, AI-2 accumulation is the limiting step for the transition from the LCD to HCD QS mode, and for driving the transition from biofilm growth to biofilm dispersal. Although CAI-1 inhibits the CqsS kinase when only a few thousand *V. cholerae* cells/mL are present, the CAI-1-CqsS circuit functions via QS, not self sensing, as released CAI-1 is shared between producing and non-producing cells (Fig 7).

**Fig 8.**
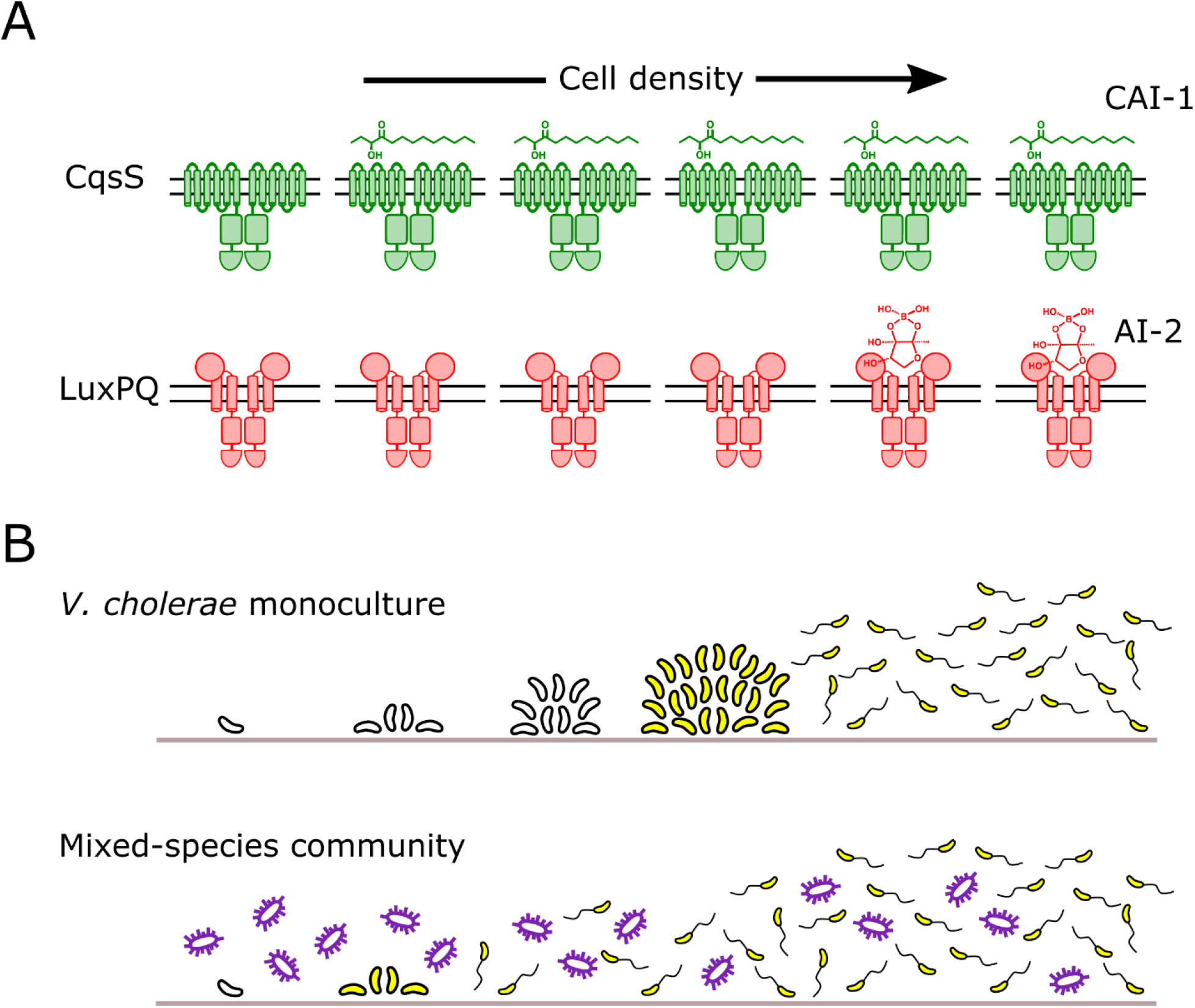
Asymmetric autoinducer thresholds drive distinct intra-genus and inter-species QS responses. (A) CAI-1 produced by *V. cholerae* engages its cognate CqsS receptor at very low cell densities. In contrast, AI-2 does not accumulate to sufficient levels to engage its cognate LuxPQ receptor until much higher cell densities. (B) The consequence of asymmetric receptor occupancy coupled with the QS system functioning as a coincidence detector is that AI-2 sets the pace at which QS occurs. In *V. cholerae* monoculture (top), the absence of AI-2 at low cell density is required for biofilm formation. Thus, exogenous AI-2, such as that provided in mixed-species communities by bacteria that possess LuxS, presumably represses *V. cholerae* biofilm development (bottom).

A longstanding mystery in the *vibrio* QS field is how the kin (CAI-1) and non-kin (AI-2) autoinducers are decoded given that they feed information into the same regulatory network. A central question has been whether each autoinducer can uniquely modulate gene expression. The present work gives us the first clues concerning this issue. The coincidence detector property of the QS system, coupled with the dramatic difference in the cell-density-dependent activation thresholds for the two autoinducers, provides a mechanism for each autoinducer to drive unique behaviors. In so doing, each autoinducer can play a fundamentally different role in the progression from LCD to HCD QS behavior (Fig 8B). Specifically, the CAI-1/CqsS circuit has a remarkably low threshold for cell-density-dependent activation. Thus, we propose that the CAI-1/CqsS arm serves as a filter that prevents the transition to HCD mode when fewer than the critical threshold number of kin cells are present, even in scenarios in which dense populations of non-kin bacteria are present (as judged by AI-2 levels). By contrast, the AI-2/LuxPQ circuit has a high cell-density dependent activation threshold, and thus, for *V. cholerae*, the buildup of AI-2 is the rate limiting step for satisfying the coincidence detector constraint. We propose that the accumulation of AI-2 sets the pace of *V. cholerae* QS. Our finding that endogenous production of AI-2 by *V. cholerae* does not exceed the threshold for LuxPQ activation until millions of cells/mL are present provides *V. cholerae* the capacity to tune into exogenous sources of AI-2, however, only after the requirement for the presence of CAI-1 is met. In our experiments, we supplied the AI-2 stimulus, but in natural contexts, exogenous AI-2 would be provided by other, non-kin bacteria in mixed-species communities.

Our demonstration that WT *V. cholerae* is sensitive to AI-2 but not to CAI-1 at all cell densities above a few thousand cells/mL indicates that when *V. cholerae* cell density has exceeded the CAI-1/CqsS activation threshold, the appearance of AI-2 would drive dramatic changes in gene expression (Fig 8B). We take this finding to mean that when a minority *V. cholerae* community of kin detects a majority of non-kin AI-2 producers, *V. cholerae* disperses from biofilms, exiting the current locale, presumably to identify superior territory. Indeed, our results further suggest that *V. cholerae* would only begin forming a new biofilm when it locates an unoccupied new area to colonize, as judged by the absence of autoinducers.

Intriguingly, the dominance of the LuxPQ receptor over the CqsS receptor in establishing the LCD QS program that we discover here has not been observed in a murine model of *V. cholerae* infection [17]. In infant mice, QS receptor kinase activity is required for colonization to occur. Mutant *V. cholerae* strains containing only the CqsS/CAI-1 or only the LuxPQ/AI-2 circuit can both establish infections. In the context of our current work, the finding that is particularly surprising is that the mutant possessing only the CAI-1/CqsS circuit is capable of colonization given the propensity of the CqsS receptor to transition from kinase to phosphatase. We suspect that in this model mammalian host, perhaps CAI-1 is degraded, a host factor sequesters CAI-1, fluid flow in the gut removes CAI-1, or reduced CAI-1 production occurs in the host. Any of these mechanisms, or others, would result in CqsS acting as a kinase to maintain the LCD QS state, and drive biofilm formation and virulence gene expression, which are required for infection.

In contrast to what we find here, in which exogenous AI-2 is the strongest QS signal, previous studies, including from us, have reported that CAI-1 is the stronger of the two autoinducers in promoting the *V. cholerae* HCD QS mode [10,11,39]. These earlier conclusions were based on data from Δ*cqsA* and Δ*luxS* mutants that produce no CAI-1 or no AI-2, respectively. We now know that positive feedback on *cqsS* transcription occurs at HCD while there is no evidence for feedback on *luxPQ* [39]. Indeed, the left panel of Fig 5B shows the cell-density-dependent increase that occurs in CqsS-3XFLAG relative to LuxQ-3XFLAG. This regulatory arrangement leads to increased CqsS levels relative to LuxPQ levels at HCD, abrogating the coincidence detection requirement. Apparently, QS coincidence detection is relevant only at cell densities below the threshold for activation of the CqsS positive feedback loop (feedback occurs at ∼OD_600_ > 1). Perhaps, once the cell density condition is reached for positive-feedback on *cqsS, V. cholerae* is at sufficiently high cell numbers that it commits to the planktonic lifestyle irrespective of the level of AI-2 in the vicinal community.

Collectively, this work, for the first time, reveals the constraints enabling self and non-self QS signaling to occur in *V. cholerae*. Although both QS autoinducers work in concert, *V. cholerae* relies on a census of the total bacteria in the local community, as measured by AI-2 concentration, to inform its decision to disperse from biofilms. For AI-2 to properly function as an inter-species signal, it is critical that kin community members do not saturate their AI-2 receptors with endogenously produced AI-2. Our work shows that *V. cholerae* avoids this circumstance by having a low cell density threshold for activation by the self, CAI-1, molecule and a dramatically higher cell-density threshold for activation by the non-self, AI-2, molecule. We predict that other bacterial species that release and detect “universal” signals must employ analogous mechanisms to prevent tripping of their QS circuits absent an accurate estimation of the total cell density of the environment.

## Materials and methods

### Bacterial strains and reagents

The parent *V. cholerae* strain used in this study was WT O1 El Tor biotype C6706str2 [43]. Antibiotics, when necessary, were used at the following concentrations: ampicillin, 100 μg/mL; kanamycin 50 μg/mL; polymyxin B, 50 μg/mL; streptomycin, 500 μg/mL; spectinomycin, 200 μg/mL, and chloramphenicol, 1 μg/mL. Strains were propagated in lysogeny broth (LB) supplemented with 1.5% agar or in liquid LB with shaking at 30° C. All strains used in this work are reported in S1 Table.

### DNA manipulation and strain construction

To generate DNA fragments used in natural transformations, including fusions and exchanges of *luxPQ* and *cqsS* alleles, splicing overlap extension PCR was performed using iProof polymerase (Bio-Rad) to combine DNA pieces. In all cases, approximately 3 kb of upstream and downstream flanking regions, generated by PCR from *V. cholerae* genomic DNA were included to ensure high chromosomal integration frequency. DNA fragments that were not native to *V. cholerae* were synthesized as g-blocks (IDT) or were purchased as plasmids (mNG was licensed from Allele Biotech) [44,45]. HapR was fused to mNG as previously described [9].

All *V. cholerae* strains constructed in this work were generated by replacing genomic DNA with DNA introduced by natural transformation (MuGENT) as recently described [46,47]. Briefly, the parent strain was grown overnight from a single colony at 30° C in liquid LB medium with agitation. The overnight culture was diluted 1:1000 into fresh medium and the strain was grown to ODø00 ∼10. Cells were pelleted at 13,000 rpm in a microcentrifuge for 1 min and were resuspended at the original volume in 1x Instant Ocean (IO) Sea Salts (7 g/L). A 100 μL aliquot of this cell suspension was added to 900 μL of a chitin (Alfa Aesar) IO mixture (8 g/L chitin), and incubated overnight without agitation at 30° C. The next day, the DNA fragment containing the desired chromosomal alteration, and an antibiotic resistance cassette for integration at the neutral locus *vc1807*, were added to the cell-chitin preparation. This mixture was incubated for 12-24 h at 30° C without shaking, after which, excess IO was removed and replaced with liquid LB. The sample was vigorously shaken to remove *V. cholerae* cells from the chitin particles, and the preparation was dispensed onto LB plates containing relevant antibiotics followed by incubation at 30° C overnight. Resulting colonies were re-streaked three times on LB plates with appropriate antibiotics, after which, PCR and sequencing were used to verify correct integration of the introduced DNA fragments. Genomic DNA from these recombinant strains was used as a template for PCR to generate DNA fragments for future co-transformation, when necessary. Antibiotic resistance cassettes linked to Δ*vc1807* were a gift from Ankur Dalia.

### Real-time biofilm development and dispersal assay

Single *V. cholerae* colonies were grown overnight in a 96-well plate in 200 μL of LB medium with shaking at 30° C covered with a breathe-easy membrane (Diversified Biotech). The cultures were diluted 1:200 into fresh LB and subsequently grown for 7 h at 30° C to OD_600_ ∼2.0. The cultures were diluted to an OD_600_ of 1 x 10^-5^, a roughly a 1:200,000 dilution in M9 medium containing glucose and casamino acids (1x M9 salts, 100 μM CaCl2, 2 mM MgSO4, 0.5% dextrose, 0.5% casamino acids). These cultures were dispensed onto No. 1.5 glass coverslip bottomed 96-well plates (MatTek) and cells were allowed to attach for 1 h at 30° C. Wells were washed to remove unattached cells by removing 200 μL of medium with a multichannel pipette and replacing with 200 μL of fresh M9 medium. After three washes, 200 μL of M9 medium was added to each well and cultures were placed in a temperature-controlled chamber for microscopy (OKO labs) at 30° C. Image acquisition was initiated 1 h later.

### Exogenous administration of synthetic autoinducers and agonists

Chemical syntheses of CAI-1, the AI-2 precursor, 4,5-dihydroxy-2,3-pentanedione (DPD), and the CqsS agonist CAI-1* have been previously described [11,48,49]. Each compound was added to medium at a final concentration of 5 μM resulting in a final DMSO concentration of 0.25%. Control cultures were supplemented with 0.25% DMSO. For experiments involving AI-2, the medium was supplemented with 0.1 mM boric acid. In all cases, autoinducers were added to cells post-attachment to glass coverslips, immediately after the final washing step described above.

### Microscopy and image analysis

Imaging of growing and dispersing biofilms was performed using a DMI8 Leica SP-8 point scanning confocal microscope. The light source for both fluorescence and brightfield microscopy was a tunable white-light laser (Leica; model # WLL2; excitation window = 470–670 nm). Biofilms were imaged using a 10X air objective (Leica, HC PL FLUOTAR; NA: 0.30) or a 63X water immersion objective (Leica, HC PL APO CS2; NA: 1.20) as indicated. For both transmission brightfield and confocal fluorescence microscopy, many wells in each plate were imaged simultaneously as specified in the Leica LasX software with a time interval of 30 min. The focal plane was maintained with adaptive focus control. A depth of 40 μm was sectioned with Nyquist sampling in XY and Z at each timepoint. Brightfield images were acquired at 640 nm and light was detected in the transmitted path using a brightfield PMT for the Leica DMI stand. For fluorescence microscopy, excitation wavelengths of 503 and 558 nm were used for mNeonGreen and mRuby3, respectively. Sequential line scanning was performed to minimize spectral bleed-through in images. Emitted light was detected using GaAsP spectral detectors (Leica, HyD SP) and timed gate detection was employed to minimize the background signal.

Image analysis was performed in FIJI software (Version 1.52e). Biofilms were segmented in the brightfield images using an intensity threshold after image smoothing. The same threshold was applied to all images in this study. The total amount of light attenuated within each segmented area was summed for the entire imaging field at each timepoint, akin to a local optical density measurement. Data were exported for quantitation and graphing in R software using gglplot2 (https://ggplot2.tidyverse.org). In all plots, data were normalized to the reference strain/conditions. In the case of fluorescence images, biofilms were initially segmented using the brightfield approach described above. Total fluorescence signal from mNeonGreen and mRuby3 was subsequently measured from single biofilms and plotted in ggplot2.

### Bioluminescence assay

Three colonies of each strain to be analyzed were individually grown overnight in 200 μL LB with shaking at 30° C in a 96-well plate covered with a breathe-easy membrane. The following morning, the cultures were diluted 1:5000 into fresh SOC medium or SOC medium containing the indicated concentration of autoinducers. The plates were placed in a BioTek Synergy Neo2 Multi-Mode reader with constant shaking at 30° C. Both OD_600_ and bioluminescence from the chromosomally integrated *lux* operon were measured. Results were exported to R, and bioluminescence values were divided by OD_600_ to produce relative light units (RLU). Results from the triplicate experiments were averaged and plotted using the ggplot2 plugin for R.

### Western Blotting

Cultures of strains carrying CqsS-3xFLAG and LuxQ-3xFLAG were collected at the indicated OD_600_ and subjected to centrifugation for 2 min at 13,000 rpm. The pellets were flash frozen, thawed and lysed for 10 min at 25° C by resuspending in 75 μL Bug Buster (Novagen, #70584–4) supplemented with 0.5% Triton-X, 50 μl/mL lysozyme, 25 U/mL benzonase nuclease, and 1 mM phenylmethylsulfonyl fluoride (PMSF) per 1.0 OD_600_ of pelleted culture. The cell lysate was solubilized in 1x SDS-PAGE buffer for 1 h at 37° C. Samples were loaded onto 4–20% Mini-Protein TGX gels (Bio-Rad). Electrophoresis was carried out at 200 V until the loading buffer reached the bottom of the gel. Proteins were transferred from the gels to PVDF membranes (Bio-Rad) for 1 h at 4° C at 100 V in 25 mM Tris buffer, 190 mM glycine, 20% methanol. Membranes were blocked for 1 h in PBST (137 mM NaCl, 2.7 mM KCl, 8 mM Na2HPO4, 2 mM KH2PO4, and 0.1% Tween) with 5% milk, followed by three washes with PBST. Subsequently, membranes were incubated for 1 h with a monoclonal Anti-FLAG-Peroxidase antibody (Sigma) at a 1:5,000 dilution in PBST with 5% milk. After washing four times with PBST for 10 min each, membranes were exposed using the Amersham ECL western blotting detection reagent (GE Healthcare). For the RpoA loading control, the same general protocol was followed except that the primary antibody was Anti*-Escherichia coli* RNA Polymerase α (Biolegend) used at a 1:10,000 dilution and the secondary antibody was an Anti-Mouse IgG HRP conjugate antibody (Promega) also used at a 1:10,000 dilution.

### Flow cytometry analyses

The secrete-and-sense strains constitutively produced mRuby3 enabling differentiation from the non-fluorescent sense-only strains. In all cases, strains were grown overnight with shaking at 30° C, either in monoculture, or as 1:1 co-cultures of secrete-and sense and sense-only strains. The cultures were diluted to OD_600_ of 5 × 10^-6^, a roughly a 1:500,000 dilution. Starting 2 h post inoculation, aliquots of cells were collected in 1 h intervals and fixation was performed per safety protocol for performing flow cytometry with a BSL2 organism. Cells were pelleted in a microcentrifuge at 13,000 rpm for 1 min, resuspended in 100 μL of 3.7% formaldehyde (Electron Microcopy Sciences) in filter-sterilized PBS, and left at room temperature for 10 min. Subsequently, three washes were performed to remove excess formaldehyde. In the three washing steps, the cells were pelleted in a microcentrifuge at 13,000 rpm for 1 min and resuspended in 1 mL of PBS. After the final wash, cells were resuspended in 1 mL of PBS, except for LCD cultures, which were resuspended at 5X concentration in 200 μL PBS to increase the frequency of detection events in the subsequent flow cytometry analysis. Following fixation and washing, cells were stored at 4° C in the dark until flow cytometry was performed. mRuby3 and mNG fluorescence signals were compared before and after fixation by microscopy and no fluorescence signal was lost during fixation.

Flow cytometry was performed on samples using a FACSAria Special Order Research Product driven by FACSDiva software (BD Biosciences). A 561 nm laser line was used to excite mRuby3 fluorescence and a 488 nm laser line was used for mNG fluorescence. Forward and side-scatter were used to gate a distinct single cell population, and within this gate, two distinct peaks were identified in the mRuby3 channel corresponding to cells that strongly produced the mRuby3 fluorescent protein (secrete-and-sense cells), and those that did not (sense-only cells). Cells were further gated based on this histogram to assign appropriate mNG signals to the secrete-and-sense and sense-only cell populations. Data from all samples were collected with identical gates, laser intensity, and PMT voltages.

---

**S1 Fig.**
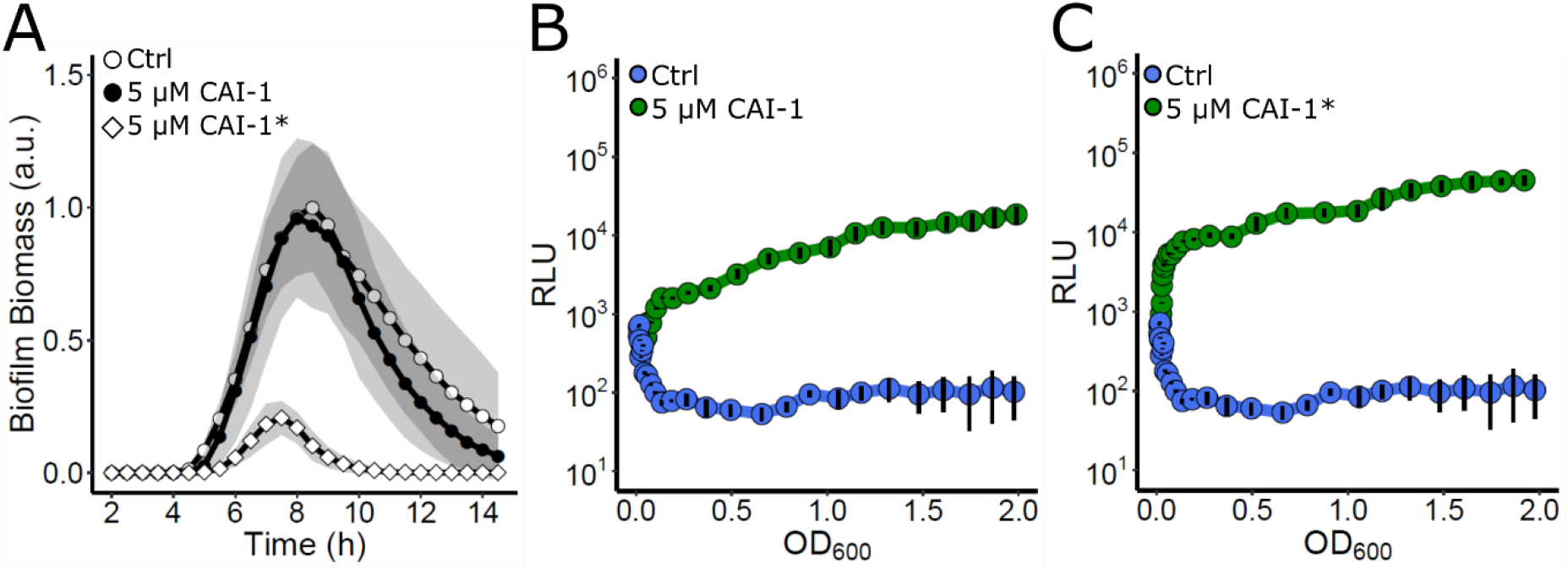
Response of the *V. cholerae* CAI-1 reporter strain to exogenous CAI-1 and CAI-1*. (A) Quantitation of biofilm biomass for the *V. cholerae* CAI-1 reporter strain (Δ*vpsS*, Δ*cqsR*, Δ*luxQ*, Δ*cqsA*) treated with 0.25% DMSO (Ctrl), 5 μM CAI-1, or 5 μM CAI-1*, over time. Data are represented as means normalized to the peak biofilm biomass of the DMSO control strain. n=3 biological and n=3 technical replicates, ± SD (shaded). (B) The corresponding *lux* pattern for the strain in A following treatment with 0.25% DMSO (Ctrl) or 5 μM CAI-1. (C) As in B following treatment with 0.25% DMSO (Ctrl) or 5 μM CAI-1*. Relative light units (RLU) are defined as light production (a.u.) divided by OD_600_. For B and C, n=3 biological replicates and error bars represent SD.

**S2 Fig.**
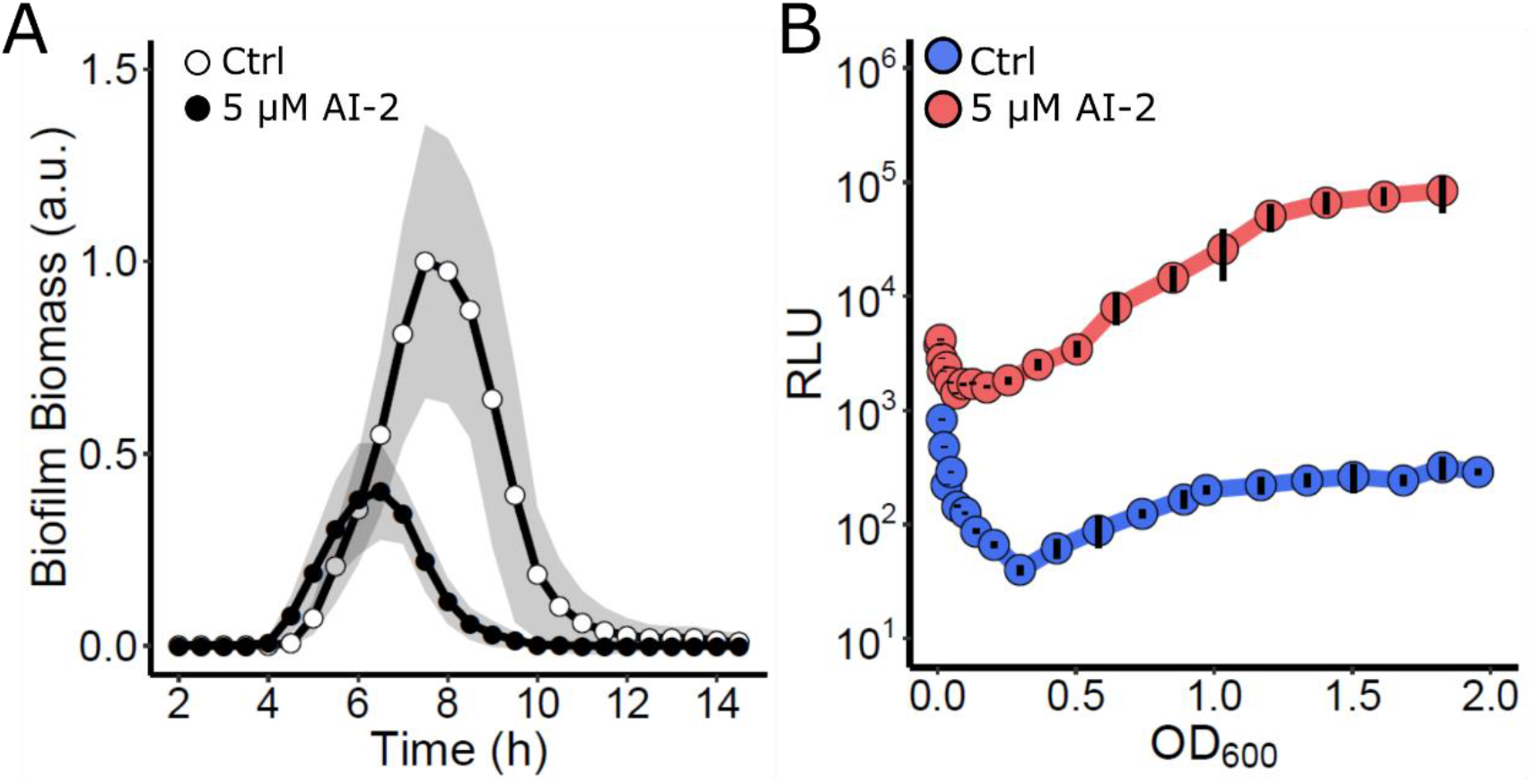
Response of the *V. cholerae* AI-2 reporter strain to exogenous AI-2. (A) Quantitation of biofilm biomass for the *V. cholerae* AI-2 reporter strain (Δ*vpsS*, Δ*cqsR*, Δ*cqsS*, Δ*luxS*) treated with 0.25% DMSO (Ctrl) or 5 μM AI-2 over time. Data are represented as means normalized to the peak biofilm biomass of the DMSO control strain. n=3 biological and n=3 technical replicates, ± SD (shaded). (B) The corresponding *lux* pattern for the strain in A following treatment with 0.25% DMSO (Ctrl) or 5 μM AI-2. Relative light units (RLU) are defined as light production (a.u.) divided by OD_600_. n=3 biological replicates and error bars represent SD.

**S3 Fig.**
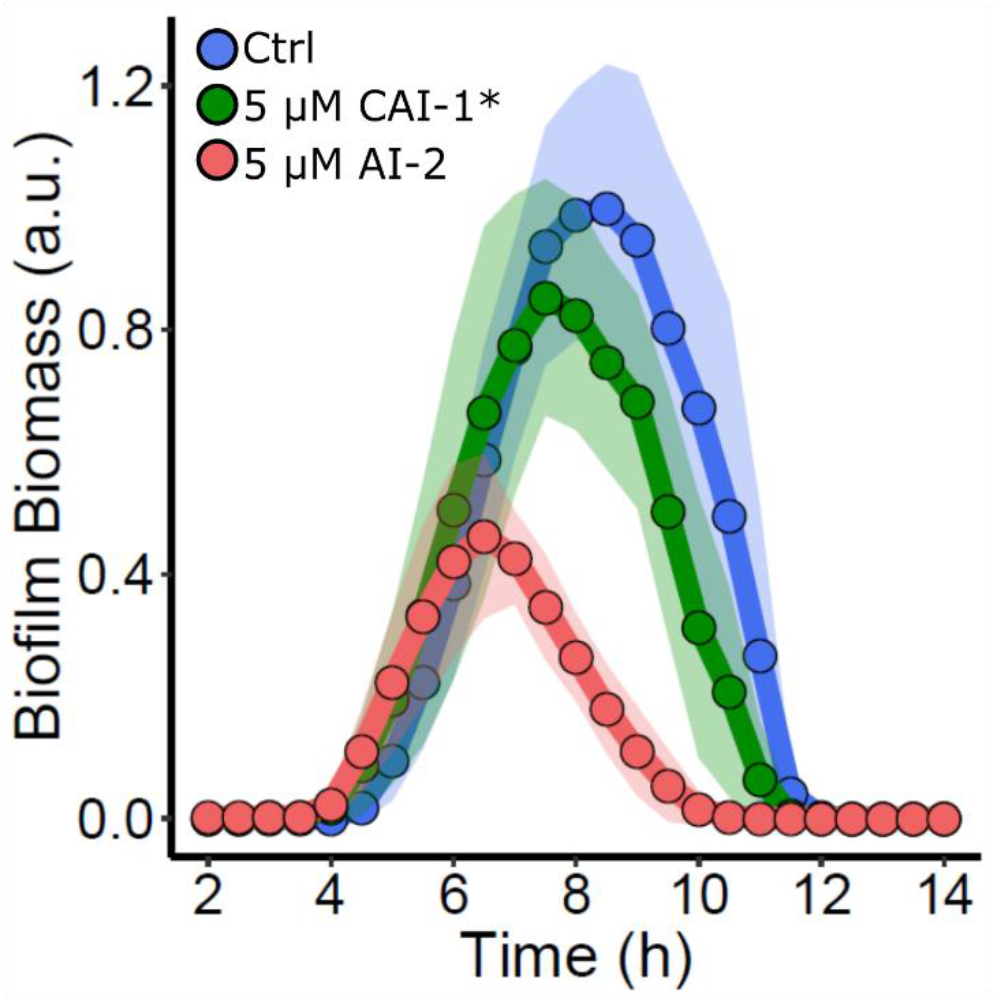
Exogenous AI-2 but not CAI-1* represses biofilm formation in the Δ*vpsS*, Δ*cqsR V. cholerae* strain. Quantitation of biofilm biomass for the *V. cholerae* Δ*vpsS*, Δ*cqsR* strain treated with 0.25% DMSO (Ctrl), 5 μM CAI-1*, or 5 μM AI-2 over time. Data are represented as means normalized to the peak biofilm biomass of the DMSO control strain in each experiment. n=3 biological and n=3 technical replicates, ± SD (shaded).

**S4 Fig.**
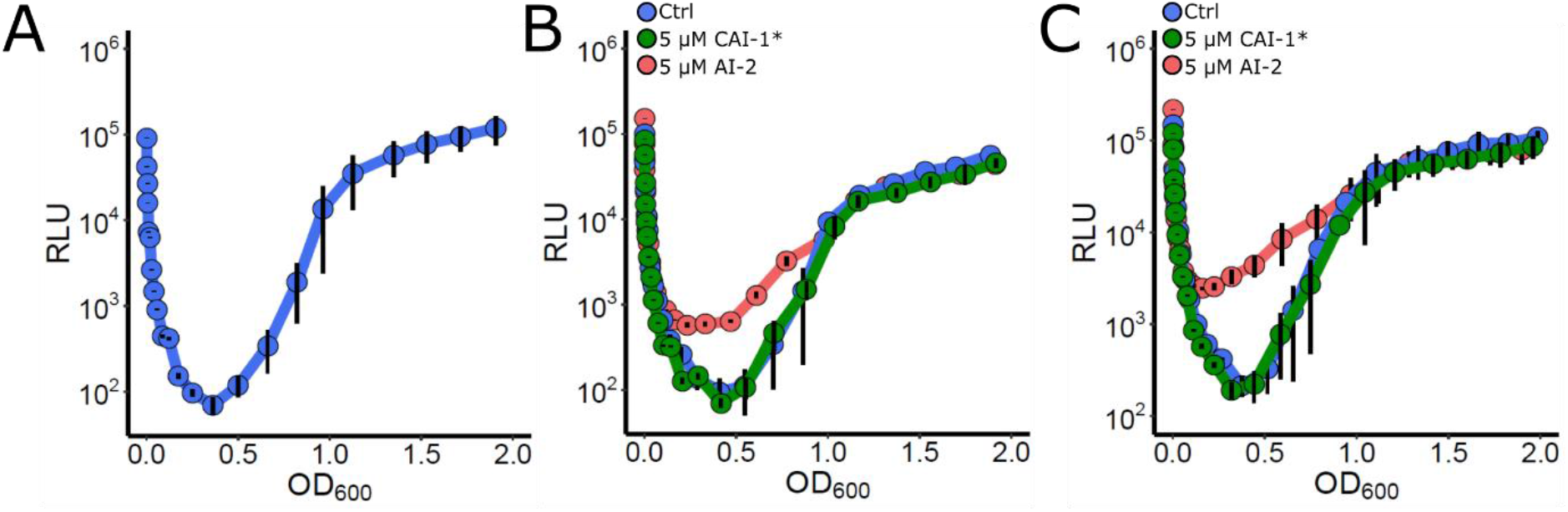
Exogenous AI-2 but not CAI-1* activates WT *V. cholerae lux* expression. (A) The *lux* pattern for WT *V. cholerae* over time. (B) As in A following treatment with 0.25% DMSO (Ctrl), 5 μM CAI-1*, or 5 μM AI-2. (C) As in B for the Δ*vpsS*, Δ*cqsR* strain. Relative light units (RLU) are defined as light production (a.u.) divided by OD_600_. n=3 biological replicates and error bars represent SD.

**S1 Table.**
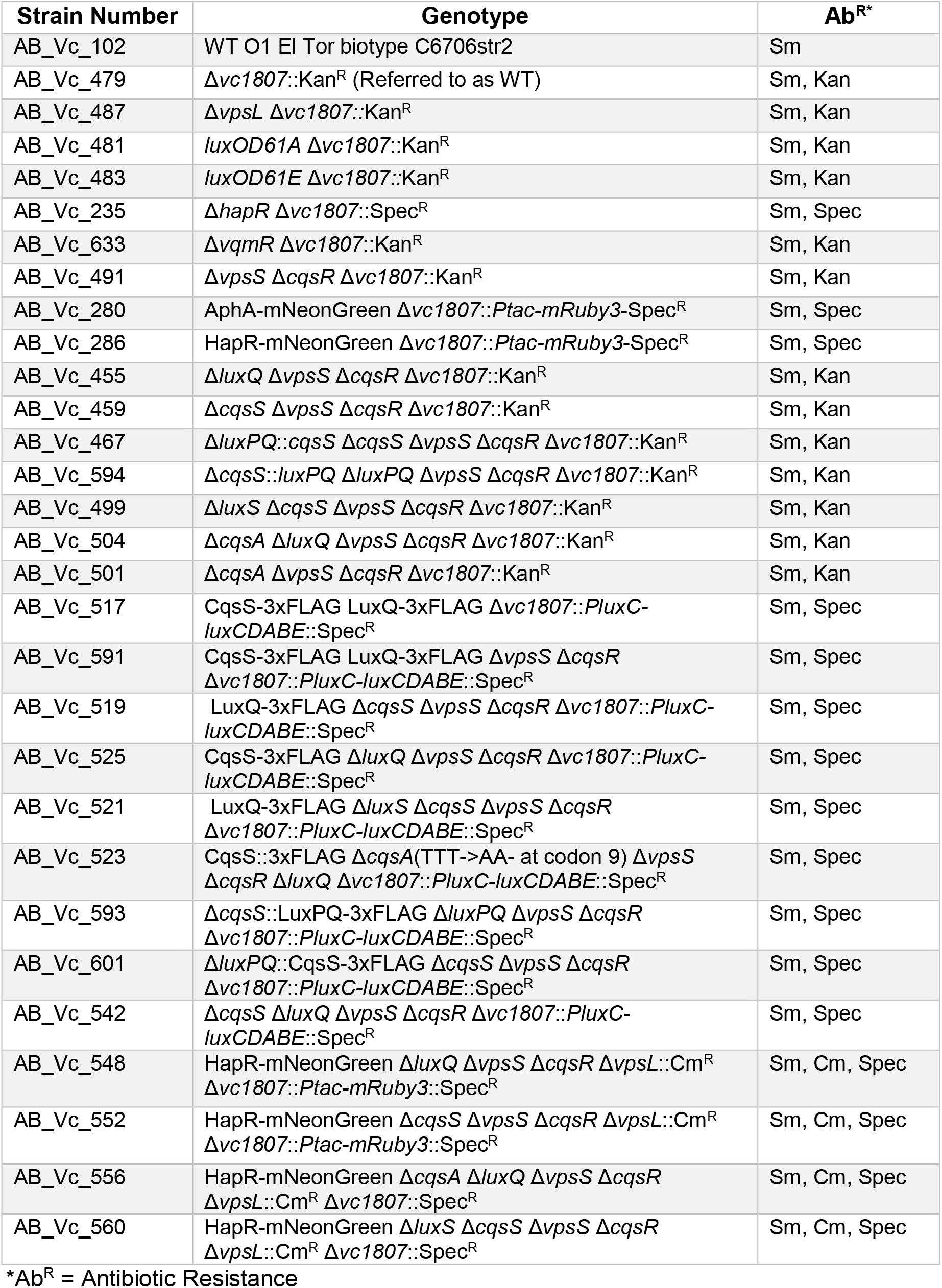

## Acknowledgements

We thank members of the Bassler group and Prof. Ned Wingreen for thoughtful discussions. We particularly thank Dr. Ameya Mashruwala for providing the luciferase reporter used in this study. This work was supported by the Howard Hughes Medical Institute, NIH Grant 5R37GM065859, National Science foundation Grant MCB-1713731, and a Max Planck-Alexander von Humboldt research award to B.L.B. A.A.B. is a Howard Hughes Medical Institute Fellow of the Damon Runyon Cancer Research Foundation, DRG-2302-17.

